# A synthetic lethal dependency on casein kinase 2 in response to replication-perturbing drugs in RB1-deficient ovarian and breast cancer cells

**DOI:** 10.1101/2022.11.14.516369

**Authors:** Daria Bulanova, Yevhen Akimov, Wojciech Senkowski, Jaana Oikkonen, Laura Gall-Mas, Sanna Timonen, Manar Elmadani, Johanna Hynninen, Sampsa Hautaniemi, Tero Aittokallio, Krister Wennerberg

## Abstract

Treatment of patients with high-grade serous ovarian carcinoma (HGSOC) and triple-negative breast cancer (TNBC) includes platinum-based drugs, gemcitabine, and PARP inhibitors. However, resistance to these therapies develops in most cases, highlighting the need for novel therapeutic approaches and biomarkers to guide the optimal treatment choice. Using a CRISPR loss-of-function screen for carboplatin sensitizers in the HGSOC cell line OVCAR8, we identified *CSNK2A2*, the gene encoding for the alpha’ (α’) catalytic subunit of casein kinase 2 (CK2). Expanding on this finding, we confirmed that the CK2 inhibitors silmitasertib and SGC-CK2-1 sensitized many, but not all, TNBC and HGSOC cell lines to the drugs that perturb DNA replication, including platinum drugs, gemcitabine, and PARP inhibitors. We identified RB1 tumor suppressor deficiency as a prerequisite context for the CK2 inhibition-mediated sensitization to these therapeutics. In RB1-deficient cells, CK2 inhibition resulted in accumulation of cells in S phase of the cell cycle, associated with micronuclei formation, and accelerated PARP inhibitor-induced aneuploidy and mitotic cell death. Patient HGSOC organoids that lacked RB1 expression displayed an enhanced long-term response to carboplatin and PARP inhibitor niraparib when combined with silmitasertib, suggesting RB1-stratified efficacy in patients. As RB1 deficiency affects up to 25% of HGSOC and 40% of TNBC cases, CK2 inhibition, proven safe from previous clinical exploration with silmitasertib, is a promising approach to overcome resistance to standard therapeutics in large strata of patients.

## Introduction

High-grade serous ovarian carcinoma (HGSOC) and triple-negative breast cancer (TNBC) are malignancies with low patient survival due to the frequent development of therapy resistance^1,2^. Standard-of-care therapy for both cancer types includes DNA-damaging agents, such as platinum drugs (alone or in combination with taxanes) or gemcitabine. Due to similar molecular characteristics (high genomic instability, high prevalence of *BRCA* mutations, and functional homologous recombination (HR) deficiency)^3^, PARP inhibitors have recently been approved for the treatment of HGSOC and TNBC. Maintenance therapy with PARP inhibitors extends the progression-free survival of patients with *BRCA* mutant tumors^4,5^. However, despite the initial success of the stratified approach, patients in this “better prognosis” cohort still often develop resistance to both chemotherapy and PARP inhibitors^6–8^, leading to disease relapses. This urges the development of strategies to improve therapeutic responses in HGSOC and TNBC patients.

Effective cancer treatments often rely on synergistically acting therapeutic agents. A synergy is defined as a cytotoxic effect of independent pharmacological perturbations that significantly exceeds the cancer cells’ viability reduction caused by either of the perturbations alone, and it can be assessed using methods like BLISS, Loewe, HSA and others^9,10^. The extreme case of a synergistic interaction is a synthetic lethality, where the combination of two non-lethal perturbations together becomes fully lethal (such that 0+0=1). In cancer therapy, this is typically applied in cases where one cancer-specific genetic alteration causes a *de novo* dependence on a separate targetable protein or cellular signal^9,10^. The therapeutic advantage of these types of interactions are that they become cancer cell specific, while normal non-malignant cells are unaffected by the drug addition. The synthetic lethal interaction between *BRCA* tumor suppressor gene mutations and PARP inhibition is an encouraging example of targetable synthetic lethality that led to effective, stratified, and even curative treatments for ovarian and breast cancer patients^2,11^. Indeed, advances in high-throughput chemical biology or functional genomics (CRISPR-based loss- or gain-of-function screens) streamline the search for synergistic and synthetic lethal combinations as druggable, context-specific targets for combinatorial cancer therapy.

Our CRISPR loss-of-function screening and validation studies identified CK2 kinase catalytic subunit α’ as one of the essential factors for carboplatin treatment survival in HGSOC models deficient for *RB1* tumor suppressor gene. Further, we demonstrate that CK2 inhibition selectively sensitizes RB1 protein-deficient HGSOC cells to carboplatin, gemcitabine, and PARP inhibitors. This is the first evidence of a combinatorial benefit of concurrent CK2 and PARP inhibition. The combination of PARP inhibitor niraparib and CK2 inhibitor silmitasertib induced aneuploidy and mitotic cell death in RB1-deficient HGSOC cells. Combinations of silmitasertib with either carboplatin or niraparib induced cytotoxicity in RB1-deficient patient-derived HGSOC organoid cultures. Similarly, we observed the synergy between silmitasertib and carboplatin or niraparib in RB1-deficient TNBC cell lines. Given that *RB1* tumor suppressor deficiency due to genomic or expression loss occurs in 15-25% of HGSOC^12–14^ and 20-40% of TNBC cases^3,15^, silmitasertib appears to be a potential partner drug to boost the therapy efficacy in such patients. Co-inhibition of CK2 and PARP provides a promising strategy to broaden the cohort of HGSOC and TNBC patients benefiting from a genetically stratified treatment to cases with RB1 deficiency. Our study provides a rationale for RB1 deficiency-guided clinical treatment selection to stratify patients in clinical trials and, ultimately, to achieve higher efficacy of the therapy in triple-negative breast and high-grade ovarian cancer.

## Results

### Genome-wide CRISPR screen identifies novel modulators of carboplatin sensitivity

To identify genes whose ablation increases the carboplatin sensitivity in HGSOC cells, we performed a pooled genome-wide CRISPR loss-of-function screen (Figure 1A) in the OVCAR8 cell line, which carries mutant *TP53*, (ubiquitous in HGSOC^12,16^) and displays functional loss of HR-mediated DNA repair^17^.

**Figure 1.**
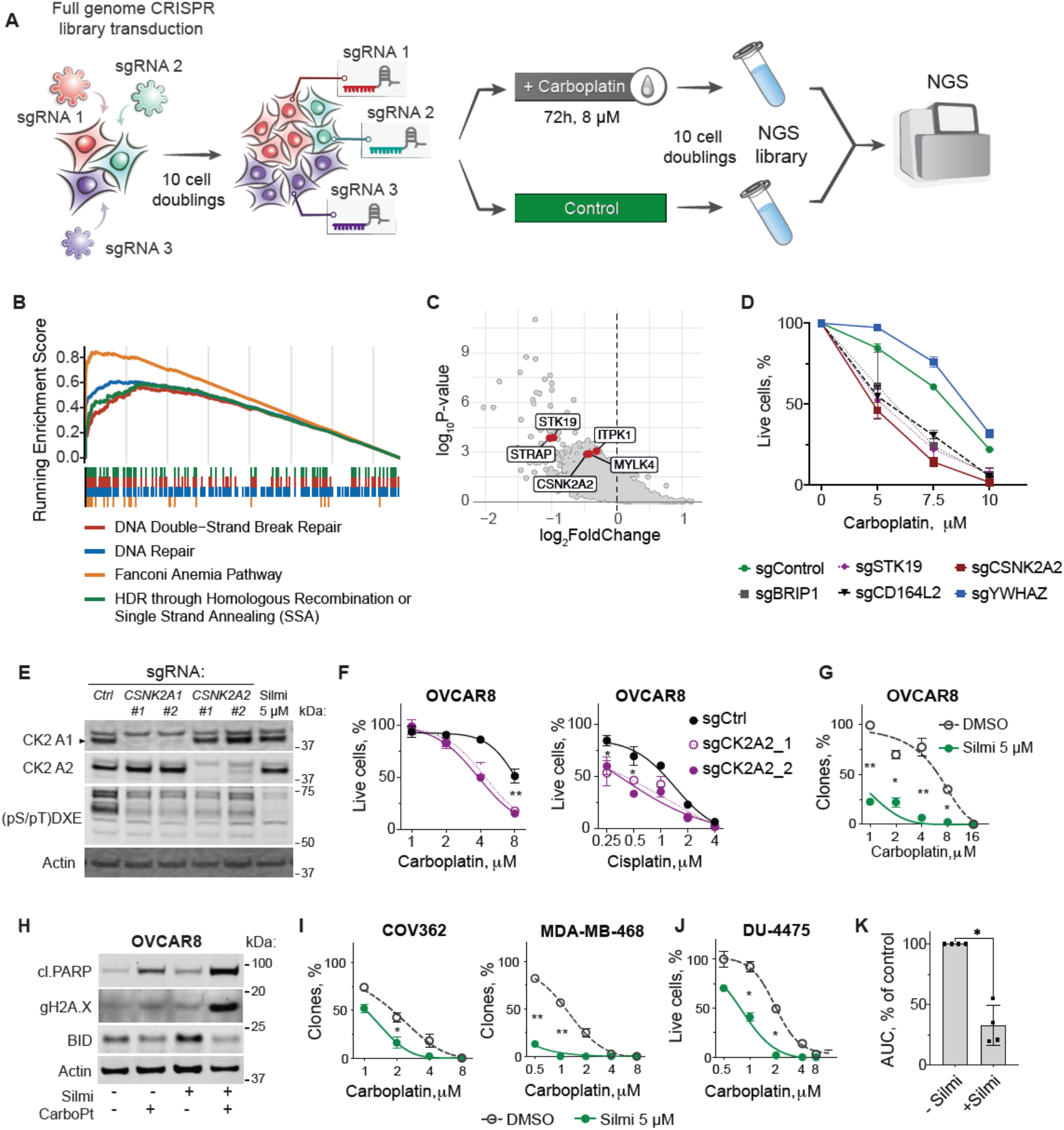
CRISPR-Cas9 knock-out screen identifies CK2 subunit α’ among novel carboplatin-sensitizing factors. **A**, Schematic representation of the CRISPR-Cas9 screen experiment. Upon selection with puromycin, the pools were united and propagated for 10 divisions to allow the elimination of the cells with fitness gene knockouts. **B**, Gene Ontology analyses conducted using Enrichr^18^ reveals significant enrichment in DNA repair factors among genes essential for carboplatin resistance. **C**, Volcano plot, the hit genes encoding for kinases are highlighted. **D**, Viability of OVCAR8 cells with the knockouts of screening hit genes upon carboplatin treatment. The numbers of live and dead cells were determined using image cytometry and Hoechst / CellTox green staining, n=2, data points represent mean±SD. **E**, Immunoblotting for CK2 kinase catalytic ubunits and phosphorylation motif in OVCAR8 cells with genetic ablation of CSNK2A1/2 or treated with CK2 inhibitor silmitasertib 5 µM (Silmi) for 4 h. **F**, Viability of OVCAR8 cells with CK2 a’ subunit CRISPR knock-out. The cells were incubated with the indicated concentrations of platinum agents for 7 days. G, Quantification of the clonogenic survival assay for indicated cell lines treated with carboplatin in the presence of 5 μM silmitasertib or vehicle for 5 days and then grown in drug-free conditions for 9 more days. The data points represent mean±SD, n=2, N=2 for each cell line. **H**, immunoblotting for apoptotic markers in VCAR8 cells treated for 5 days with 5 μM silmitasertib, 8 μM carboplatin, or their combination. **I**, Quantification of the clonogenic survival assay, performed as in G. **J**, Quantification of the survival assay for DU-4475 cells treated with carboplatin in the presence of 5 μM silmitasertib or vehicle for 5 days and then grown in drug-free conditions for 9 more days. Due to growth in suspension, live cells were counted using Trypan blue exclusion assay and Countess II cell counter. The data points represent mean±SD, n=2, N=2. **K**, Difference in the area under the curve (AUC) for carboplatin response curves in OVCAR8, COV362, MDA-MB-468, DU-4475 in the presence of silmitasertib. \**P* < 0.05; \*\**P* < 0.01, \*\*\**P* < 0.001, for D, F, G, I, J, two-way ANOVA with Tukey post-test, for K, Mann-Whitney test.

As expected, the analysis of the sgRNA abundances upon DNA-damaging carboplatin treatment identified multiple DNA repair factors as the most enriched genes among the statistically significant hit genes (Figure 1B, Supplementary tables ST1-2). We further focused on druggable carboplatin-sensitizing genes, such as genes encoding kinases (*CSNK2A2, STK19, ITPK1, MYLK4, STRAP*) or receptor molecules (*CD164L2, HTR5A*), that appeared among the top-100 genes ranked by MAGECK RRA score as genes that promote survival after the carboplatin treatment (Figure 1C, Supplementary Table ST1). To validate the screen results, we re-tested several hit genes with different RRA scores using individual sgRNAs. In agreement with the genome-wide CRISPR screen, the targeted knock-outs of sensitizer genes (*CSNK2A2, STK19, CD164L2*, and top-1 hit *BRIP*) or resistance genes (*YWHAZ*) reduced or improved the viability of OVCAR8 cells in response to carboplatin, respectively (Figure 1D).

We chose to explore the carboplatin-sensitizing effect of the knock-out of *CSNK2A2* encoding the α’ catalytic subunit of the serine-threonine kinase CK2, which can be targeted with selective compound silmitasertib^19,20^, and has a documented role in DNA damage response (reviewed in^21^). We modeled the loss of CK2 kinase activity by knocking out catalytic subunit-encoding genes *CSNK2A1* (α) and *CSNK2A2* (α’). Knock-outs of α and α’ subunits only partly decreased the levels of CK2 phosphorylated substrates (pS/pT)DXE, Figure 1E), suggesting that the CK2 catalytic domain consisting of a-a or a’-a’ homodimers is still capable of maintaining partial CK2 activity in OVCAR8 cells. Chemical inhibition with silmitasertib, a selective CK2 inhibitor, efficiently inhibited CK2 substrate phosphorylation levels (Figure 1E). Both genetic and chemical inhibition of CK2 activity (Figure 1E) reduced the viability of OVCAR8 cells in response to platinum drugs (Figure 1F-G). Consistent with its greater inhibitory effect on CK2-mediated phosphorylation, silmitasertib had a carboplatin-sensitizing effect both in short-term viability assays and in long-term clonogenic survival tests (Figure 1G). OVCAR8 treatment with carboplatin and silmitasertib induced apoptotic markers, PARP cleavage, γH2AX accumulation, and BID degradation (Figure 1H), indicating the cytotoxic effect of the combination.

We tested the carboplatin-sensitizing effect of silmitasertib in a broader panel of cell models. In addition to HGSOC, we tested cell lines from TNBC, as the two cancers have similar molecular signatures^3^ and treatment options (platinum drugs, gemcitabine, and PARP inhibitors)^2^. We observed a cytotoxic sensitization by silmitasertib in 5 out of 14 tested HGSOC and TNBC cell lines (Figure 1I-K and S1), which indicated a stratified effect of the combination.

### CK2 inhibition sensitizes cancer cells to PARP inhibitors and gemcitabine

Carboplatin kills cancer cells by inducing multiple types of DNA damage and cell cycle defects, including GpG intrastrand and interstrand DNA crosslinks that lead to replication fork stalling and double-strand breaks. Mechanisms of protective responses in cells include the activation of G1/S and G2/M checkpoints^6^, allowing for efficient DNA repair before DNA replication and cell division continues. To determine what types of carboplatin-induced DNA lesions or cell cycle defects cause cytotoxicity upon CK2 inhibition, we performed an imaging cytometry-based combinatorial drug screen testing the sensitizing interaction between silmitasertib and 10 other agents that induce DNA damage via different mechanisms relevant to carboplatin cytotoxicity (Figure 2A, Supplementary Table ST3). We compared the drug combination responses in three cell lines that displayed a sensitizing carboplatin-silmitasertib interaction (OVCAR8, COV362, MDA-MB-468, Figure 1G, I), and four cell lines where CK2 inhibition did not have sensitizing effect with carboplatin (OVCAR3, COV318, OVCAR4, KURAMOCHI, Figure S1A-D).

**Figure 2.**
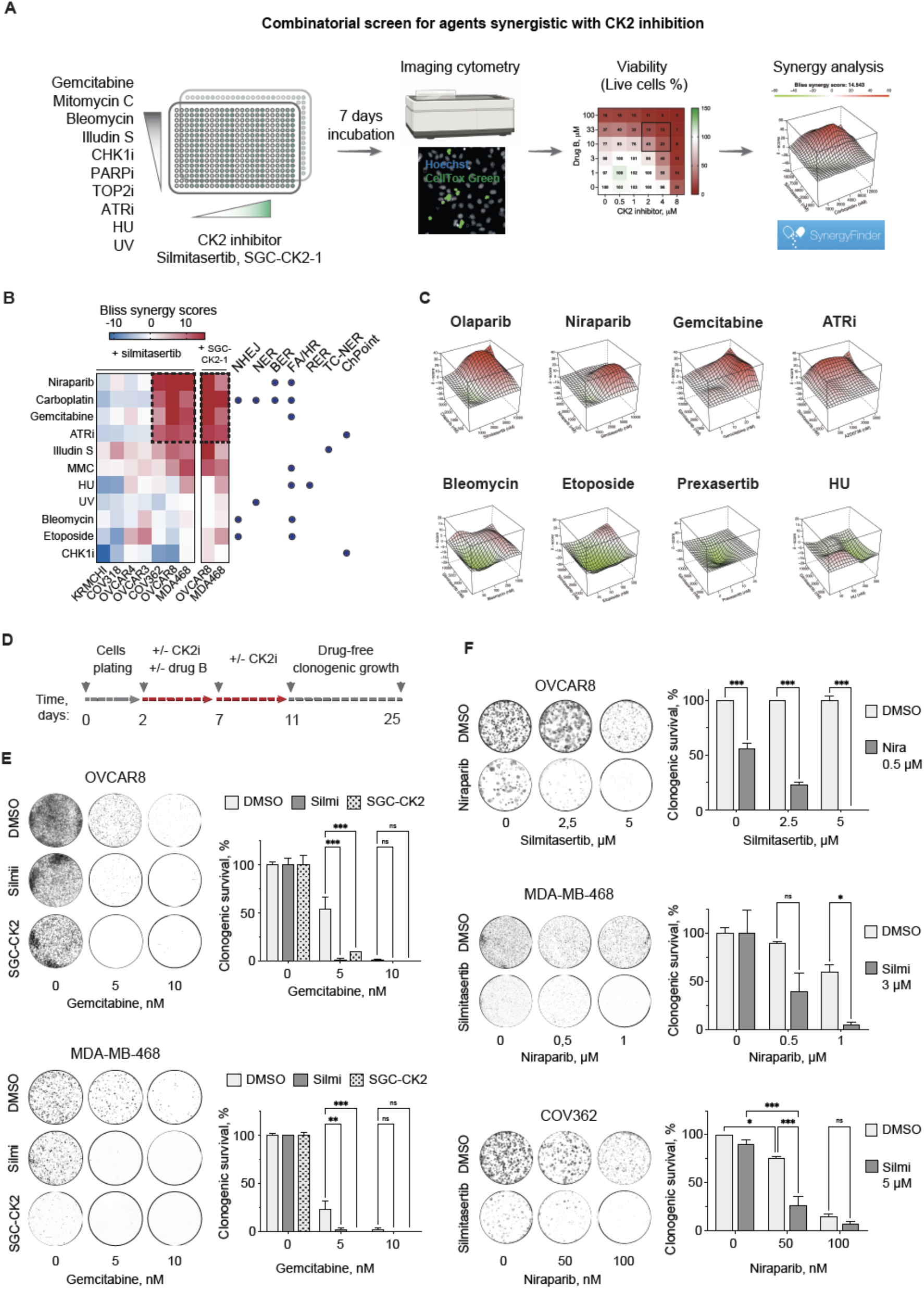
Interactions between silmitasertib and replication-perturbing drugs in HGSOC and TNBC cells. **A**, Schematic depiction of the combinatorial drug screening approach. The cell lines (OVCAR8, OVCAR3, OVCAR4, COV362, COV318, and MDA-MB-468) were plated to multi-well plates with single drugs or drug combinations pre-dispensed at a range of doses. After 7 days of incubation, the number of live and dead cells for each treatment was measured by image cytometry using Hoechst and CellTox Green dyes. Viability was calculated as a fraction of the live cells normalized to DMSO-treated controls. **B**, Heatmap representing Bliss synergy scores for drug-drug interactions in the tested cell lines (mean of 2 independent experiments for each cell line). NHEJ, non-homologous end joining; NER, nucleotide excision repair; BER, base excision repair; FA/HR, Fanconi anemia / homologous recombination repair pathway, RER, ribonucleotide excision repair; TC-NER, transcription-coupled nucleotide excision repair. The dots mark the published evidence for the contribution of the pathways to the repair of the lesions by the tested drugs. **C**, Representative 3D Bliss synergy plots for synergistic (Bliss score > 8) and additive or antagonistic (Bliss score< 4) drug-interactions in OVCAR8. **D**, Schematic timeline of the clonogenic survival test for validation of the screen results. **E**, Clonogenic survival of OVCAR8 and MDA-MB-468 treated with gemcitabine and CK2 inhibitors silmitasertib or SGC-CK2i. Bars represent the gemcitabine-surviving fraction of clones normalized to DMSO control or CK2 inhibitor-only control, respectively and show mean±SD for one representative experiment. The experiments were performed twice. **F**, Clonogenic survival of the indicated cell lines treated with niraparib and silmitasertib. The experiments were performed twice. Bars represent mean±SD for one representative experiment, n=2. \**P* < 0.05; \*\**P* < 0.01, \*\*\**P* < 0.001, two-way ANOVA with Tukey post-test.

The drug screen showed that CK2 inhibition also enhanced the toxicity of PARP inhibition, ATR inhibition and gemcitabine in OVCAR8 COV362, and MDA-MB-468 cells, but not in the other tested models (Figure 2B-C, Supplementary Figures S2-3). These agents induce replication-associated DNA damage and, according to chemogenomic profiling, the sensitization to these agents is associated with dysfunctional replication fork quality control and Fanconi anemia pathway/inter-strand crosslink (FA/ICL) repair^22^. Illudin S, which induces transcription-associated DNA damage repaired by TC-NER, and mitomycin C, which induces non-distorting crosslinks repaired by the FA pathway, showed an additive effect with silmitasertib. The lack of a sensitizing effect in the responses to illudin S and mitomycin C between cell lines suggested that these mechanisms of action were not similarly vulnerable to CK2 inhibition as replication-associated DNA damage induced by carboplatin, gemcitabine, or PARP inhibition. CK2 inhibition failed to potentiate the toxicity of double-strand break inducers etoposide and bleomycin, dNTP-depleting hydroxyurea, checkpoint kinase 1 (CHK1) inhibitor prexasertib or UV light (Figure 2B-C, Supplementary Figure S2-3). This evidence argued against the contribution of CK2 activity to cell survival upon DNA lesions repaired by end-joining, NER, RER, or CHK1-mediated checkpoint activation.

We further validated the sensitizing interaction between CK2 inhibition and PARP inhibitors or gemcitabine in a clonogenic survival test (Figure 2D-F). Silmitasertib or the more selective CK2 inhibitor SGC-CK2-1^23^ enhanced the toxicity of gemcitabine in OVCAR8 and MDA-MB-468 cells (Figure 2E). Similarly, concurrent CK2 and PARP inhibition reduced the long-term survival of HGSOC cell lines OVCAR8 and COV362 and TNBC cell lines MDA-MB-468 and DU-4475 (Figure 2F, Supplementary figure S4A-C). CRISPR-mediated knock-out of *CSNK2A2* led to a moderate, but significant decrease in the clonogenic survival of OVCAR8 cells upon niraparib treatment (Figure S4D), consistent with the drug screening results. Collectively, the results demonstrated that CK2 inhibition enhanced the toxicity of agents that induce replication-associated DNA damage in a subset of TNBC and HGSOC cell lines.

### CK2 inhibition deregulates S phase progression in RB1-deficient cells

Since CK2 inhibition potentiated the cytotoxicity of the drugs that induce replication-associated DNA damage, we hypothesized that CK2 inhibition differentially influences G1/S or S cell cycle phase progression in the sensitized cell lines. To test this, we analyzed the cell cycle progression in silmitasertib-treated cells by EdU incorporation assay. In agreement with our hypothesis, in OVCAR8 and MDA-MB-468, CK2 inhibition increased both the EdU incorporation rate and S phase subpopulation fraction in both flow cytometry and imaging assays (Figure 3A-B, 3E, S4E-F). In contrast, in OVCAR3, COV318, OVCAR4, and KURAMOCHI, silmitasertib had a cytostatic effect as reflected by a decrease both in the fraction of EdU-incorporating cells and in the intensity of EdU incorporation (Figure 3C-D, E, S4F) and a dose-dependent decrease in the mitotic index (Figure 3F). These data indicated that CK2 kinase inhibition appears to result in a failed G1/S checkpoint in the sensitized cell lines (cerulean shaded in Figure 3), while the checkpoint remains intact in the non-sensitized cell lines (violet shaded in Figure 3).

**Figure 3.**
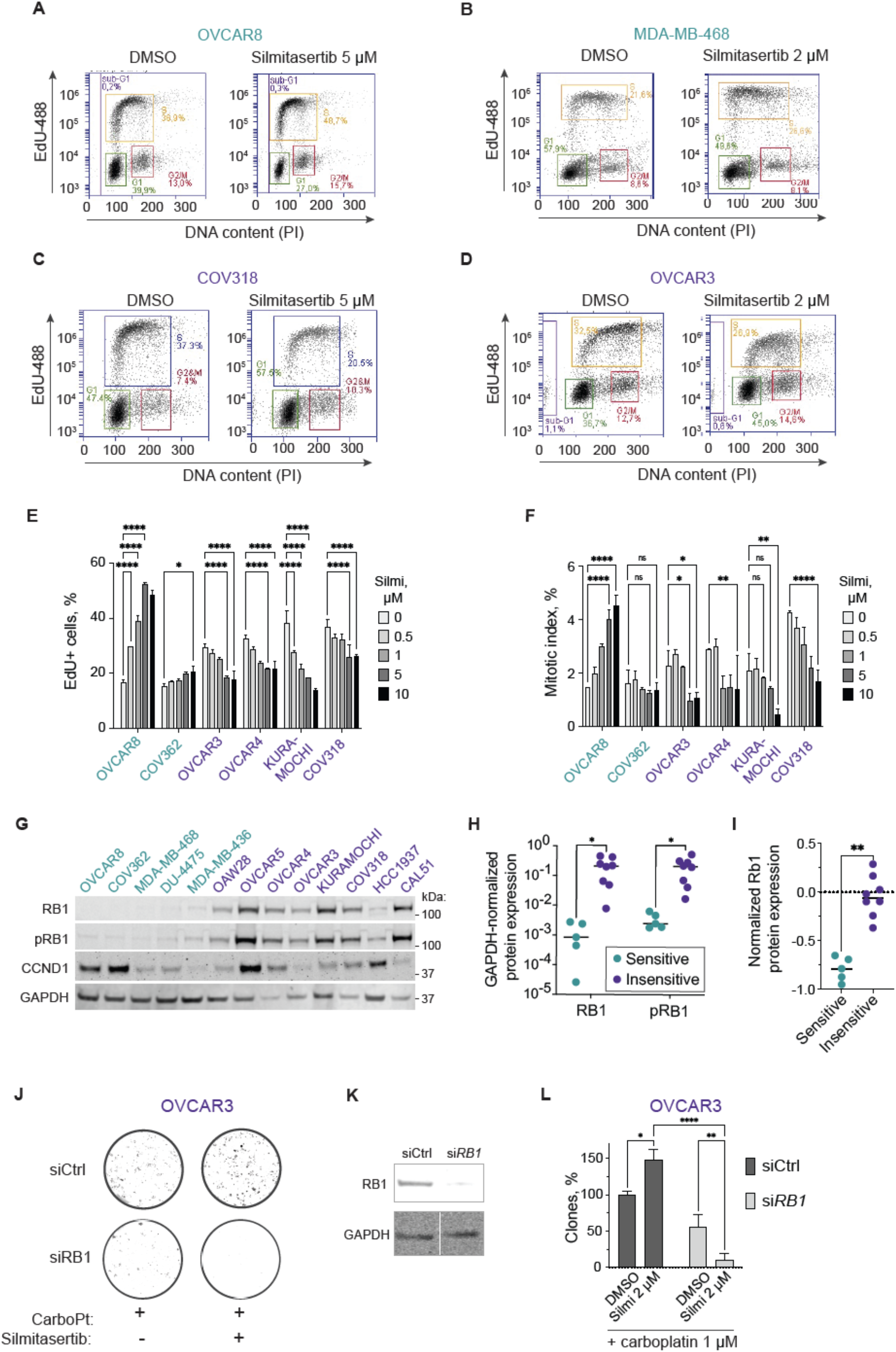
The CK2 inhibition-dependent sensitization to carboplatin or niraparib in HGSOC and TNBC cell lines depends on loss of RB1 expression. Cell lines were classified as sensitive or insensitive if they presented or did not present sensitization to carboplatin by silmitasertib in the long-term survival assay related to Figure 2, respectively. **A-D**, Flow cytometry analysis of the EdU incorporation. The cell lines were pretreated with silmitasertib or vehicle control for 72 h, pulse-labeled with 5 µM EdU in the growth media for 30 min, and immediately collected for fixation and detection of EdU using Click-iT assay. **E**, Quantification of the EdU incorporation imaging in the indicated cell lines pretreated with silmitasertib for 72 h. **F**, Mitotic index quantification. Adherently growing cells were incubated with silmitasertib for 72 h, fixed, and stained with Hoechst. Linear classifier algorithm-based detection of mitotic figures was used to assess the % of mitotic nuclei out of the total number of nuclei on the confocal microscopy images. **G**, Western blotting analysis of RB1, phospho-RB1 (T608), and Cyclin D1 expression in cell lines. **H**, Quantification of RB1 and phospho-RB1 immunoblotting in G. **I**, RPPA-based RB1 expression comparison between cell lines from CCLE study^24^. Classification of the cell lines as in H. **J**, Clonogenic survival of RB1-depleted OVCAR3 cells. 18 h post-transfection with 30 nM siRNA, the cells were treated with 1 µM carboplatin, 2 µM silmitasertib, or their combination for 48 h and replated to 6-well plates for clonogenic growth for 7 days. **K**, Western blot for RB1 expression upon si*RB1* transfection. **L**, Quantification of the clonogenic assay in J. Number of colonies in carboplatin-treated control siRNA transfectants is taken as 100%. For E, F, and L, the bars represent mean±SD, n=2, N=2. \**P* < 0.05; \*\**P* < 0.01, \*\*\**P* < 0.001, two-way ANOVA with Tukey post-test. For H and I, \**P* < 0.05; \*\**P* < 0.01, Mann-Whitney test.

To identify factors contributing to the differential cell cycle effects and the sensitization to carboplatin, PARP inhibitors, and gemcitabine by CK2 inhibition, we analyzed the expression of G1/S and S phase regulators in HGSOC and TNBC cell lines in publicly available gene expression datasets (Cancer Cell Line Encyclopedia^24^, (CCLE). We found differential expression of RB1 pathway genes stratifying the cell lines (Figure S5A), with a loss of *RB1* gene expression characteristic for the cell lines sensitive to chemotherapy combinations with silmitasertib. Protein expression analysis confirmed the loss of RB1 protein in all four cell lines where silmitasertib sensitized the responses to carboplatin and niraparib (Figure 3G-H). Publicly available reverse-phase protein array (RPPA) data on RB1 expression confirmed our western blot findings (Figure 3I). The association with CK2 inhibition mediated sensitization was not observed for cyclin D1, another commonly deregulated G1/S transition mediator (Figure 3G).

To assess whether the loss of RB1 protein could induce CK2 inhibition-mediated sensitization to carboplatin, we transfected the insensitive, RB1-proficient OVCAR3 cell line with siRNA against the *RB1* transcript, and then tested its response to carboplatin in clonogenic survival test. Even partial *RB1* depletion rendered OVCAR3 cells hypersensitive to carboplatin while the control transfected cells were partially protected by silmitasertib (Figure 3J-L, S5B-D). Altogether, our findings indicate that the loss of RB1 expression induces a dependence on CK2 in controlling the S phase progression and in response to the therapeutics that induce replication-associated DNA damage in HGSOC and TNBC models.

### CK2 inhibition exacerbates aneuploidy and mitotic cell death in niraparib-treated RB1-deficient cells

To determine if CK2 inhibition enhances niraparib toxicity in a particular cell cycle phase, we followed the cell division in RB1-deficient and -proficient cells using time-lapse microscopy. Exposure to niraparib alone for 48 h elevated the incidence of mitotic cell death and aneuploidy in all tested cell lines (Figure 4A, S6), in agreement with the previous reports^25–27^. In the presence of silmitasertib, niraparib-treated RB1-deficient cells displayed an increase both in the number of mitotic catastrophes and aneuploid divisions (Figure 4A, S6), indicating that mitotic aberrancies contribute to the increased cytotoxicity of the niraparib-silmitasertib combination. Notably, we observed extended mitosis duration even for normal divisions resulting in two daughter cells in combination-treated RB1-deficient lines (Figure 4B). The CK2 inhibition alone did not affect the mitotic outcome, however, it induced a dose-dependent accumulation of post-mitotic micronuclei in RB1-deficient, but not in RB1-proficient, cell lines (Figure 4C-D). Additionally, with observed exacerbated micronucleation in RB1-deficient cells after carboplatin treatment (Figure S4F). Together, the data suggest that CK2 plays a role in maintaining mitotic fidelity that becomes critical on the background of RB1 deficiency.

**Figure 4.**
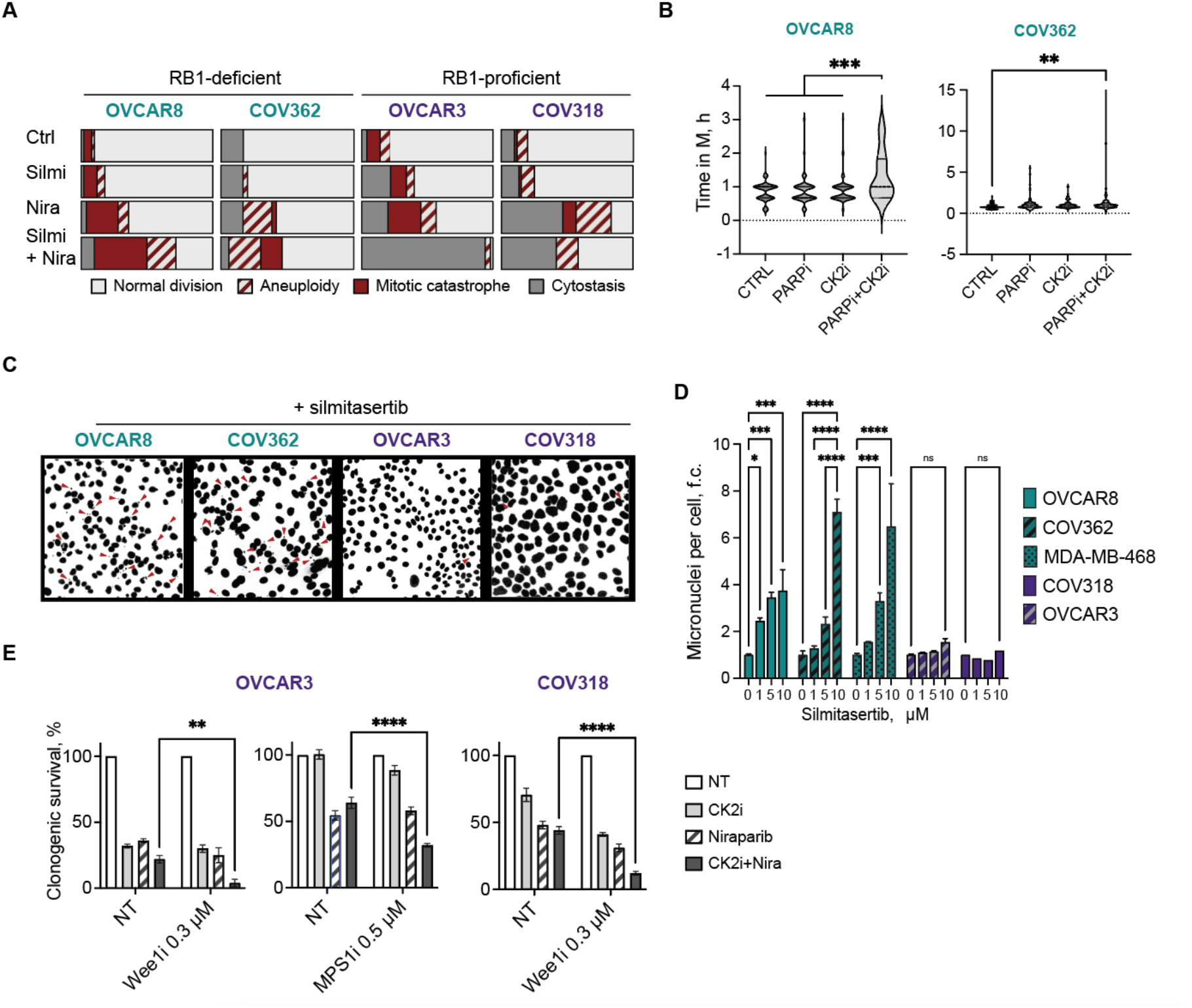
CK2 inhibition triggers mitotic cell death and post-mitotic aneuploidy in RB1-deficient HGSOC cells. **A**, Outcomes of mitoses in sensitive/RB1-deficient and insensitive/RB1-proficient HGSOC cell lines exposed to PARP inhibitor niraparib for 48 h in the presence or absence of silmitasertib. Cells were not synchronized. Cells were considered cytostatic if no mitotic rounding and division were observed during 48 h of observation. N=2, 50-60 cells were analyzed per each condition. **B**, Time from prometaphase until cytokinesis for the mitotic division resulting in 2 daughter cells in OVCAR8 and COV362. Drug treatment: niraparib for 48 h in the presence or absence of silmitasertib before the start of the time-lapse imaging (every 15 min for 96 h total). Cells were not synchronized. **C**, Representative images of Hoechst-stained nuclei of the indicated cell lines upon 120 h of treatment with silmitasertib. Red arrows indicate micronuclei. **D**, Quantification of the imaging experiment in C. At least 500 nuclei were counted for each condition. N = 2. **E**, Clonogenic survival of RB1-proficient cell lines treated with silmitasertib and niraparib for 72 h. Wee1 inhibitor adavosertib was added for the last 18 h of the drug treatment, after which the cells were released in the drug-free medium. For MPS1 inhibition, AZ3146 was added for 72 h together with CK2 and PARP inhibitors. N = 2. \**P* < 0.05; \*\**P* < 0.01, \*\*\**P* < 0.001, two-way ANOVA with Tukey post-test.

To assess the role of cell cycle checkpoints in response to the CK2 and PARP inhibitors combination in RB1-proficient cells, we tested the sensitivity in presence of cell cycle checkpoint inhibitors. Inhibition of Wee1 kinase for 18 h prior to exposure to silmitasertib-niraparib combination sensitized RB1-proficient cell lines COV318 and OVCAR3 (Figure 4E). The inhibition of spindle assembly checkpoint kinase MPS1 had a similar effect in OVCAR3 (Figure 4E). We concluded that the activity of cell cycle checkpoints in RB1-proficient cells may play a protective role against the CK2+PARP inhibitor combination at the division onset.

The post-mitotic chromatin aberrations and mitotic cell death following PARP inhibitor^28^ or platinum treatment often result from unresolved replication-born DNA lesions^29,30^. We used a common approach^31^ to address whether the PARP inhibitor sensitization by CK2 inhibition resulted from the induction of replicative DNA damage. CK2 inhibition alone did not induce nuclear γH2AX and RPA loading in S phase cells, characteristic markers of the replicative DNA damage^31^, in the tested cell lines (Figure S7A-B, Figure S7C). In addition, CK2 inhibition alone did not affect the level of phosphorylated CHK1 (pS317) in the RB1-deficient OVCAR8 cells (Figure S8A) or the formation of FANCD2 foci (Figure S8B), the marker of inter-strand crosslink repair and replication fork protection complex active in S phase^32,33^. Similarly, in the RB1-deficient OVCAR8 cells, silmitasertib treatment did not affect the FANCD2 regulatory ubiquitination reported to be CK2 activity-dependent^34^ (Figure S8A). We did not observe significant changes in RPA and γH2AX loading in response to acute replication stress inducer HU or PARP inhibitor upon silmitasertib treatment (Figure S8C), arguing against exacerbation of replicative DNA damage by CK2 inhibitor.

We also examined whether HR-mediated DNA repair counteracted the sensitization to carboplatin or niraparib by CK2 inhibition in RB1-proficient cells. However, neither BRCA2 knockdown nor inhibition of Rad51 filament formation conferred enhanced the CK2 inhibitor sensitization to PARP inhibition or affected the formation of micronuclei in response to silmitasertib in RB1-proficient cells (Supplementary Figure S9), suggesting that the sensitivity to CK2 and PARP co-inhibition did not depend on the activity of this DNA repair mechanism.

### Silmitasertib potentiates the long-term efficacy of carboplatin in patient-derived RB1-deficient HGSOC organoids

Testing drug responses in long-term HGSOC *ex vivo* 3D cultures, or organoids, is a powerful functional assay to predict the therapeutic efficacy in patients^35–37^. To evaluate the translational potential of the carboplatin-silmitasertib combination we assessed its efficacy in a set of ten previously established^38^, clinically relevant, genomically characterized HGSOC patient-derived organoid cultures derived from ascites (N=5) or omentum (N=5) and collected at different stages of the treatment (Figure 5A-B). First, we determined the RB1 protein expression in actively growing organoids and classified them to RB1-deficient and -proficient groups based on the relative RB1 protein expression (Figure 5C-D). Next, we assessed the long-term survival and re-growth of the organoids after treatment with carboplatin +/- silmitasertib. The combination of silmitasertib with carboplatin substantially attenuated the survival of RB1-deficient patient-derived organoids as compared to carboplatin alone (Figure 5E). For 3 out of 5 RB1-deficient patient samples, CK2 inhibition had carboplatin-sensitizing effect during the whole experiment timeline, while for the remaining 2 samples the effect was time-point dependent (EOC989, EOC382), reflecting sample-specific differences in carboplatin sensitivity. In contrast, silmitasertib did not enhance the cytotoxicity of the platinum agent in RB1-proficient samples, and, in one sample, it even protected the cells from the carboplatin cytotoxicity (Figure 5F).

**Figure 5.**
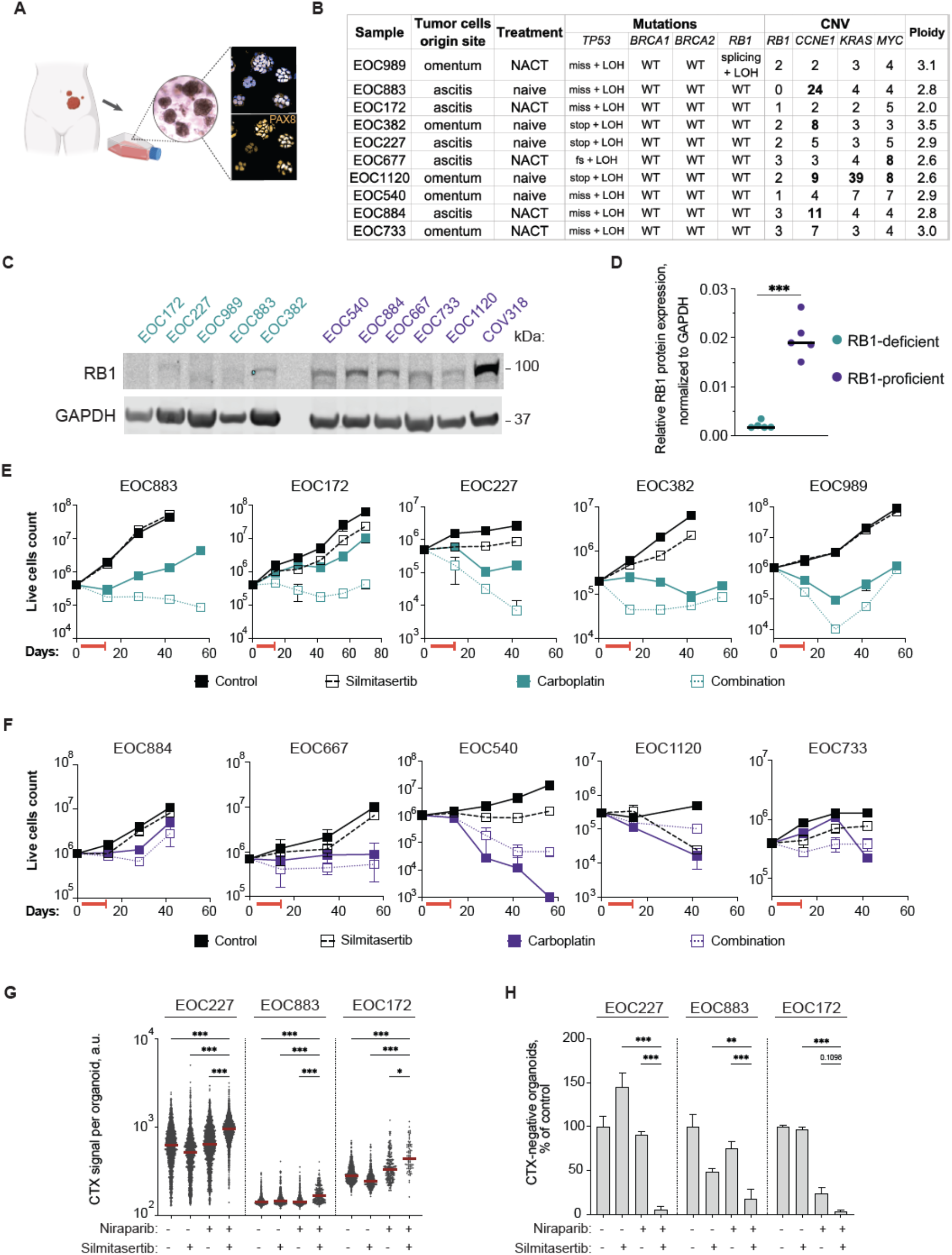
Efficacy of silmitasertib in combination with carboplatin or niraparib in HGSOC patient-derived organoids. **A**, Schematics of the establishment of the HGSOC organoids. Immunofluorescent imaging of PAX8, HGSOC histologic marker^39^, in organoids. **B**, Characteristics of the selected set of organoid. NACT, neoadjuvant platinum-based chemotherapy. Mutations: fs, frameshift; stop, stopgain; miss, missense mutation; splicing, splicing isoform mutation; LOH, loss of heterozygocity. Copy number variation columns present numbers of gene copies per cell, bolded numbers correspond to amplification of the gene. **C**, Immunoblotting analysis of RB1 protein expression in the set of organoids. **D**, Quantification of the immunoblot in C. **E, F**, Long-term survival of sensitive/RB1-deficient and insensitive/RB1-proficient organoids. The organoids were exposed to the drugs alone or in combination for 7 days, followed by additional 4 days in presence of vehicle or silmitasertib alone (red T-line, drug treatment, 11 days in total). After the drug treatment, the organoids were passaged at the density of 5×105 live cells per 200 μl of the gel for 6-8 weeks. N=2, mean±SD. **G**, Quantification of the CellTox Green fluorescent signal per organoid. N=2, N of organoids (30-300 per well) varied depending on the treatment. **H**, Viability of RB1-deficient organoids, number of CellTox Green-negative (live) organoids expressed as the % of DMSO control. Plots present mean±SD, *P < 0.05; **P < 0.01, ***P < 0.001, for C Mann-Whitney test, for G-H two-way ANOVA with Tukey post-test.

We further tested if silmitasertib potentiated the cytotoxicity of niraparib in RB1-deficient patient-derived samples EOC172, EOC227, and EOC883. Confocal imaging-based quantification of live and dead organoids discriminated by CellTox Green staining showed a significantly lower survival after treatment with niraparib in the presence of silmitasertib (Figure 5G-H, Supplementary Figure S10), indicating that CK2 inhibition enhances the efficacy of the PARP inhibition in RB1-deficient patient-derived HGSOC models. These data indicate that CK2 inhibition combined with standard therapeutics is a promising novel treatment strategy for RB1-deficient HGSOC.

## Discussion

CRISPR screening offers an effective approach to systematically identify gene-drug interaction and for identification of novel drivers of sensitivity or resistance to cancer therapeutics^40–45^. Our CRISPR screen study identified *CSNK2A2* as a carboplatin sensitizer gene, refining the previous findings on the interaction between CK2 inhibitors and platinum drugs^46–48^. Chemical inhibition of CK2 activity with two specific inhibitors confirmed the role of CK2 in carboplatin sensitivity, validating the phenotype identified in the screen. The combination of carboplatin with silmitasertib was effective in producing long-term response in models that lacked RB1 tumor suppressor protein expression. Therefore, the identified specific interaction between the drug combination and RB1 deficiency can be classified as a context-specific synthetic lethality^10^. Expanding previous findings on targetable regulators of platinum resistance^22,44,49–51^, in addition to CK2 we identified four kinases (STK19, STRAP, MYLK4, ITPK1, Figure 1, Supplementary Table ST1) that have not been implicated in carboplatin response before. Interestingly, STK19 was reported as a regulator of transcription-coupled nucleotide excision repair^22,52^, a mechanism of platinum-DNA adduct removal^53–55^. As STK19 can be targeted with the recently developed compound ZT-12-037-01^56^, our data also warrant the evaluation of the STK19 inhibitor as a carboplatin sensitizer in HGSOC models in future studies.

Our study demonstrates for the first time that, in addition to chemotherapy, CK2 inhibition enhances the efficacy of the PARP inhibitors in HGSOC and TNBC cells. The *in vitro* toxicity of the combination of CK2 and PARP inhibitors in RB1-deficient cells was associated with accumulation of S phase cells, extended mitosis, aneuploidy, and mitotic cell death, while CK2 inhibition alone triggered intensive post-mitotic aberrations - micronuclei formation. Indeed, replication stress-associated mitotic cell death and post-mitotic aberrations have been reported for cells lacking G1/S checkpoint and/or DNA repair activity^28^. However, since we did not detect an increase of DNA damage markers (γH2AX or RPA associated with chromatin) in the S phase upon CK2 inhibition, we attributed these phenotypes to the inhibition of mitotic functions of CK2 and its interaction with checkpoint kinases, such as Wee1. In favor of the latter, Wee1 inhibition sensitized RB1-proficient ovarian cancer cells to the combination of silmitasertib with carboplatin or niraparib (Figure 4E). Cancer cells with RB1 loss display higher genomic instability^57^ and have been reported to be more vulnerable to the inhibition of the regulators of the mitotic spindle assembly and cytokinesis, such as Aurora kinases, due to increased spindle assembly checkpoint activity and susceptibility to tubulin dynamics disruption^58–60^. CK2 has redundant activity with Aurora at kinetochores^61^ and, hypothetically, CK2 inhibition may lead to consequences similar to the loss of Aurora B-mediated phosphorylation, resulting in the killing of the RB1-deficient cells in the presence of accumulated DNA damage by PARP trapping or alkylation. Interestingly, CK2 catalytic subunit α’ is differentially phosphorylated in mitosis^62^, pointing at cell cycle phase-specific roles of the CK2 composed of α’-α’ catalytic core similar to Aurora B kinase activity. In addition to PARP inhibitors and platinum drugs, we found CK2 activity critical for RB1-deficient cancer cell survival after treatment with replication poisons, such as illudin S or ATR kinase inhibitors (Figure 3), suggesting a similar mechanism of toxicity for those drug combinations, while a lack of synergistic interaction with the other tested agents argues against the general sensitization to apoptosis by CK2 inhibition.

*RB1* genetic inactivation by deletions or mutations occurs in 15-25% of HGSOC^12,16^ and 20-40% of basal-like breast cancer^3^ (a gene expression–defined subtype closely corresponding to histologically-defined TNBC). Hence, the drug combination could be relevant for a substantial fraction of patients. To our knowledge, no patient stratification has been done in clinical studies of silmitasertib, either as a single agent (clinical trials NCT04663737, NCT03897036, NCT03904862, NCT01199718, NCT00891280, NCT04663737, NCT04668209) or in combination with chemotherapy (NCT02128282). Profiling RB1 deficiency in the tumors (either by DNA sequencing, transcriptomics, or immunohistochemistry) could offer a promising strategy to stratify the patients in future clinical studies of CK2 inhibitor combinatorial treatments and help retrospectively re-evaluate the outcomes of earlier studies made with unselected patient populations. Importantly, RB1 deficiency in patient-derived *ex vivo* HGSOC organoids stratifies the responses to silmitasertib combinations with platinum or niraparib. Given that tumor-derived organoids usually recapitulate the clinical responses of the tumor of origin^35,38,63,64^, the effect of CK2 inhibition on RB1-deficient organoids suggests that the respective tumors may have shown sensitivity as well. Notably, two of the “responder” organoid cultures were from the tumors of NACT-treated patients, suggesting that the combination can be of benefit for patients with platinum-resistant disease as well.

Silmitasertib is an orally administered drug candidate that has been tested in combination with platinum and gemcitabine in metastatic cholangiocarcinoma with manageable toxicity^65^. Given moderate toxicity in vivo^19,65,66^, these drug combinations may be promising for overcoming chemotherapy resistance and producing durable responses in the subset of patients with RB1-deficient tumors. The safety of CK2 inhibitor/PARP inhibitor combination has not been studied *in vivo* and needs further preclinical validation. However, the very fact that most healthy cells in the body have functional RB1, and therefore would be insensitive to the silmitasertib therapeutic combinations, argues in favor of the mild toxicity profile of the such treatment. Collectively, our data suggest that the combination of a CK2 inhibitor with platinum drugs or PARP inhibitors should be considered for clinical testing as treatment of RB1-deficient tumors. The example of CK2 inhibition potentiating the efficacy of the standard-of-care therapeutics, specifically in the context of RB1 deficiency, highlights the pressing need for better accounting for tumor molecular landscape in drug combination research, biomarker discovery, and clinical study design.

## Acknowledgements

We thank the staff of Biomedicum Helsinki Flow Cytometry Unit, and High-Content Analysis Unit, High-Throughput Biomedicine Unit, and Genomics Unit at the Institute for Molecular Medicine Finland (FIMM) for providing research infrastructure and the excellent technical help. We thank HERCULES consortium for providing access to molecularly characterized HGSOC patient samples. We are thankful to Prof. Claus Sørensen and his team (University of Copenhagen) for fruitful discussions that shaped the experimental design and helped data interpretation.

Figure 5A was created with BioRender.

## Authors’ contributions

Conception and design: D.B., Y.A., K.W. Development of methodology: D.B., Y.A., W.S., T.A., K.W. Acquisition of clinical samples and data: J.H., W.S. Acquisition of data: D.B., Y.A., L.G.M., S.T., M.E.,

W.S. Analysis and interpretation of data: D.B., Y.A., J.O. Supervision: S.H., T.A., K.W. Resources and funding acquisition: D.B., Y.A., W.S., S.H., T.A., K.W. Writing—original draft: D.B. Writing—review and editing: D.B., Y.A., W.S., J.O., M.E., S.H., T.A., K.W.

## Data availability

All data needed to evaluate the conclusions in the paper are present in the paper and/or the Supplementary Materials.

## Materials and methods

### Cell lines

For the list of cell lines and culture conditions please see Supplementary Table ST4. The cell lines were maintained in the appropriate culture medium at 37°C with 5% CO_2_ in a humidified incubator. The identities of the cell lines were confirmed using the GenePrint 10 System (Promega) at FIMM Sequencing service. Cells were routinely tested for mycoplasma negativity using MycoAlert kit (Lonza, # 11600271).

### CRISPR library production

LentiCas9-Blast vector was a gift from Feng Zhang (Addgene #52962). Human Brunello CRISPR knockout pooled library was a gift from David Root and John Doench (Addgene #73178). The library was transformed into electrocompetent Lucigen Endura™ *Escherichia coli* (Lucigen; cat. 60242-2) using a Bio-Rad MicroPulser Electroporator (#1652100), program EC1 according to the manufacturer’s protocol. The electroporated bacteria were plated onto ten 15-cm LB agar plates with 100 µg/mL ampicillin. After an overnight incubation at 32°C, the DNA plasmid was extracted using a NucleoBond® Xtra Midi kit (MACHEREY-NAGEL #740410.50). The efficiency of transformation was controlled by plating 0.01% of the transformation reaction to a 15 cm LB agar plate with 100 µg/mL ampicillin.

### Virus production

The HEK 293-FT cells were seeded at ∼ 10^5^ cells per cm^2^ 16 h prior to transfection. Transfection of lenti-Guide-Puro plasmid with CRISPR library, packaging plasmids VSV-G and psPAX2 (Addgene, #14888 and #12260) was done using Lipofectamine 2000 transfection reagent according to the manufacturer’s protocol. Viral supernatant was collected 48 h post-transfection and the titer was assessed according to Stewart et al.^*67*^.

### CRISPR screening

Cas9-expressing OVCAR8 cells were transduced with Brunello lentiviral knockout sgRNA library (4 sgRNAs/gene)^68^ at a multiplicity of infection (MOI) of 0.3. After puromycin selection, the transduced cells were propagated for 10 divisions to allow for the elimination of fitness genes-targeting sgRNAs. Next, the cell pool was divided to control and treatment pools; carboplatin at 8 μM was applied to the treatment pool for 72 h, then the drug was washed off, and both control and carboplatin-treated pools were expanded for 10 cell doublings. 8 million cells were collected 3 days after infection (T0), 10 divisions after infection (T1), and 10 divisions after carboplatin treatment started (T2) for gDNA extraction and sgRNA amplification for sequencing.

### Next generation sequencing

For CRISPR screening, the genomic DNA was extracted using NucleoSpin® Tissue kit (MACHEREY-NAGEL; #740952.50) according to the manufacturer’s protocol. sgRNA cassettes were amplified from 2.5 µg of the template in 50-μL reactions using OneTaq® DNA Polymerase (NEB; #M0480) and LG.Lib.ampl1.F and LG.Lib.ampl1.R primers (Supplementary Table ST5). Illumina sequencing primer binding sites were added by PCR amplification of sgRNA amplicons with primer mix WS Stager Mix and LG.gRNA.Ampl.NGS.R (Supplementary Table ST5). Illumina indices and adapters for sample multiplexing were added by PCR amplification with Illumina_indX_F and Illumina_indX_R primers using NEBNext® Ultra™ II Q5® Master Mix (NEB, #M0544). Samples were purified using AMPure XP beads (Beckman Coulter; #A63880). The library was sequenced with a NextSeq500 Illumina sequencer using PE100 protocol (with 10% PhiX spike-in).

For genomic profiling of patient-derived HGSOC organoids, the NGS, mutation and CNV calling was performed earlier and described by Senkowski and co-authors^38^.

### CRISPR screen analysis

Samples were demultiplexed using Illumina bcl2fastq to generate FASTQ files. Individual sgRNA counts were extracted using the *count_spacers*.*py* script^69^. Positively selected genes were identified using the MAGeCK tool^70^ and DESeq2(Wald)^71^ using simplified routines provided by DEBRA R package^72^. The over-representation and gene set enrichment analysis for GO-BP (biological process) and GO-CC (cellular component) terms were performed with clusterProfiler R package^73^ using top 150 genes with the following parameters *pAdjustMethod = “BH”, pvalueCutoff = 0*.*25, qvalueCutoff = 0*.*25*.

### Clonogenic survival assay

Cells (3-5×10^4^ per well) were plated in 6-well plates in duplicates for each treatment. After 24 h, the drugs at the indicated concentrations were added and the cells were incubated for 5 days, followed by additional growth in drug-free medium or in presence of CK2 inhibitor or vehicle for 3 days. After that, the cells were trypsinized and 10, 25, or 50% of the sample were re-plated to the fresh drug-free medium in new plates for clonogenic growth for 10 more days, during which the medium was replenished twice. For the quantification, cells were fixed with ice-cold methanol:acetic acid mix (7:1) and stained with 0.1% crystal violet. The brightfield whole-well images of the wells were taken using Cytation5 imager (BioTek) and analyzed automatically using Gen5 software (BioTek) to count the number of clones per well.

### Drug sensitivity screening

Drugs diluted in DMSO or water to the desired concentrations were dispensed at 30 nL volume to 384-well black plates (Corning, # 3864) using an Echo 550 acoustic liquid handler (Labcyte). The compounds were plated at five different concentrations in 3-fold dilutions covering a concentration range relevant for each drug. Cell-killing benzethonium chloride (BzCl, 100 μM) and compound vehicle (DMSO, 0.1%) were used as positive and negative controls, respectively. Cells were diluted to medium at the desired number per mL and the suspension was dispensed to the pre-drugged plates at 30 μL using a MultiFlo dispenser (BioTek). After 7 days of incubation at 37°C, 10 µL of PBS containing 4 μg/mL Hoechst and 1/10000 CellTox Green Dye (Promega) were were dispensed per well 1 h prior to imaging at Cytation5 image cytometer or Opera Phenix (Perkin Elmer) confocal screening microscope.

### Organoid cultures

Previously established long-term HGSOC organoid cultures^38^. were expanded in BME-2 Cultrex gel for at least 2 passages prior to drug sensitivity testing to get actively proliferating cultures. The media, passaging, dissociation were done as described^38^.

### Drug sensitivity testing on organoids

For long-term drug survival assay, dissociated organoid cells (5×10^5^-10^6^) were seeded to BME-2 gel (Cultrex, Bio-Techne) at the density of 2.5-5×10^3^ cells/µL and allowed to take up for 4 days in presence of 5 µM ROCK kinase inhibitor. After ROCKi was removed, the organoids were exposed to carboplatin or silmitasertib alone or in combination for 7 days, followed by additional 4 days in presense of vehicle or silmitasertib alone. After drug treatment, the organoids were dissociated, live cells were counted using Trypan blue exclusion method and Countess II cell counter (Invitrogen). Live cells (3-6×10^5^ or all surviving cells) were passaged to fresh BME-2 droplets at the same density. The passaging and live cell counting was repeated in a similar manner bi-weekly for 6-8 more weeks. The experiment was performed in duplicates.

For short-term drug sensitivity assay, dissociated organoid cells (2-5×10^4^) were seeded in 10 µL BME-2 gel droplets (Cultrex, Bio-Techne), one droplet per well in 96-well Cell Carrier Ultra plates (Perkin Elmer) and allowed to take up for 4 days in media containing 5 µM ROCK kinase inhibitor. Next, the medium was changed to 200 µl of medium containing niraparib, silmitasertib, or the combination. Medium and drugs were replenished after 4 days. After 7 days since drug addition, the medium was changed to drug-free, and after another 7 days of incubation at 37°C, 20 µL of PBS containing 10 μg/mL Hoechst and 1/4000 CellTox Green Dye (Promega) were dispensed per well 8 h prior to imaging at Opera Phenix (Perkin Elmer) confocal screening microscope. Image analysis was performed on the maximum projection image of 10 whole-well z-planes of the confocal imaging using Harmony software (Perkin Elmer).

### Drug combination analysis

To assess the effect of the drug combination treatments, we applied the Bliss independence model^74^. The negative values of the Bliss synergy score indicate antagonism,values around zero indicate additivity, and the positive values indicate synergy. Bliss scores were calculated for each drug combination based on the 4×4 dose-response matrix, and the Bliss synergy score values in the most synergistic area were calculated by SynergyFinder 2.0^75^, with values higher than 8 considered as robust synergy.

### Gene editing

Parental Cas9 cell lines were generated by viral infection with lentiCas9-Blast (Addgene, # 52962) followed by blasticidin selection for 7 days. sgRNAs for targeting CSNK2A1, CSNK2A2, and RB1 (Supplementary Table ST6) were cloned to lentiGuide-Puro vector (Addgene, #52963) and delivered by lentiviral transduction to the Cas9-expressing cells followed by puromycin selection (1 µg/mL) for 5 days. The efficiency of the knockouts was validated by immunoblotting.

### Lentiviral transduction

Cells were incubated with lentiviral supernatant for 24 h in the presence of 8 μg/μl of polybrene (Sigma-Aldrich) at a multiplicity of infection >5 unless otherwise stated.

### RNAi

RB1-targeting siRNA (SI00007091) and AllStarNegative control siRNA (QIAGEN) were incubated in OPTI-MEM with Lipofectamin RNAiMAX (Thermofisher) according to manufacturer’s instructions and added to the cells at the final concentration of 20 nM. After 24 h, the medium was changed and the transfectants were subjected to the treatments as stated in the respective figure legends.

### Cell cycle analysis

EdU incorporation assay was done using Click-IT kit (Thermofisher, #C10337) according to the product manual. Cells were labeled with 5 µM EdU for 30 min. DNA staining was done with propidium iodide in presence of RNAse A (100 µg/mL) for 1 h at 37°C. At least 10 000 events per sample were analyzed on an Accuri C6 Plus (Becton Dickinson). Data was analyzed using Accuri C6 Plus Sampler software.

### Immunostaining

Primary and secondary antibodies used for immunostaining are listed in Supplementary Table ST7. Cells were pre-extracted with 0.35% TX-100 in PBS on ice, fixed with 4% PFA, and after 3 washes with PBS incubated in the blocking buffer (PBS, 0.5% BSA, 20 mM glycine, 0.05% TX-100) for 1 h at room temperature. Immunostaining was performed overnight at +4°C. After 3 washes with the blocking buffer, the secondary antibodies were applied for 1 h. After washing off the secondary antibodies, counterstaining with Hoechst 33342 was done at 10 µg/mL in PBS for 5 min. The cells were imaged at an Opera Phenix confocal screening microscope (Perkin Elmer) with the 40x water immersion objective.

### Immunoblotting

Cells were lysed in RIPA buffer without EDTA supplemented with Pierce protease and phosphatase inhibitors cocktail (Thermofisher) and 20 U/mL benzonase (Millipore) on ice for 30 min. The samples were centrifuged for 15 min at 17 000 rcf at 4°C, the supernatants were processed for loading to Bis-Tris 4-12% gradient Bolt PAGs using Bolt loading dye solution and reducing agents and electrophoresis in MES running buffer (all from Thermofisher) according to the manufacturer’s instructions. Transfer of the nitrocellulose membranes was done overnight in Towbin transfer buffer, after which the membranes were stained with Ponceau Red to detect the protein transfer efficiency, washed with TBS-0.05% Tween-20 (TBS-T), and blocked in 5% non-fat milk in TBS-T. The membranes were incubated with primary antibodies listed in the Supplementary Table ST8 at the indicated dilutions in 5% non-fat milk in TBS-T overnight at +4°C except for GAPDH and beta-actin antibodies (1 h at RT). After 3 washes in TBS-T, the membranes were incubated with the secondary fluorophore-conjugated antibodies. Primary and secondary antibodies and their working dilutions used for immunoblotting are listed in Supplementary Table ST7. The membranes were scanned at LiCOR Odyssey imager and the bands’ fluorescence intensity was quantified in ImageLite software. Immunoblotting experiments were repeated twice. Representative images are shown.

### Statistics and reproducibility

Statistical analyses were conducted using GraphPad Prism (v.7.0.2). For non-parametric comparison of 2 groups of data, Mann-Whitney test was applied. For multiple comparisons, statistical significance (adjusted *P* values) was calculated using the two-way analysis of variance (ANOVA), with Tukey multiple comparisons test. Results are reported as non-significant at P > 0.05, and with increasing degrees of significance: *0.01 < *P* ≤ 0.05, **0.001 < *P* ≤ 0.01, ***0.0001 < *P* ≤ 0.001 and \*\*\*\**P* ≤ 0.0001.

