## Supplementary Figures S1-S10 for "A synthetic lethal dependency on casein kinase 2 in response to replication-perturbing drugs in RB1-deficient ovarian and breast cancer cells"

Bulanova et al.

Supplementary File 1

#### Supplementary figures legends

**Supplementary Figure S1.** A-I, Quantification of the clonogenic tests in OVCAR3, COV318, OVCAR4, KURAMOCHI, OAW28, OVCAR5, MDA-MB-436, HCC1937, CAL51 cell lines treated for 5 days with carboplatin in presence or absence of silmitasertib, followed by 3 days in silmitasertib-containing or drug-free medium, respectively, amd by another 6 days in drug-free medium for all wells. Bars represent mean±SD for one experiment, n=2, N=2. *P* values are calculated using a two-way ANOVA with Tukey post-test for multiple comparisons. *$0.01 < P \leq 0.05$, **$0.001 < P \leq 0.01$.

**Supplementary Figure S2.** Surface heat maps related to Figure 2C representing the synergy scores (Z axis) between silmitasertib (2.5, 5, 7.5 µM, Y axis) and the indicated drugs in the range of concentrations (X axis), n=3, N=2.

**Supplementary Figure S3.** A, Surface heat maps for the synergy scores (Z axis) between SGC-CK2-1 (0.3, 1, 3 µM, Y axis) and the indicated drugs in the range of concentrations (X axis). B, Surface heat maps for the synergy scores (Z axis) between silmitasertib (2.5, 5, 7.5 µM, Y axis) and paclitaxel in the range of concentrations (X axis), n=3, N=2.

**Supplementary Figure S4**. A, C, Quantification of the clonogenic survival of OVCAR8 and MDA-MB-468 treated with niraparib and CK2 inhibitor SGC-CK2-1 for 5 days. Data points represent mean ±SD, n=2, N=1 for each cell line. B, Quantification of the survival of DU-4475 TNBC cell line treated with the drugs for 5 days and allowed to grow for 9 more days in drug-free medium. Viability is calculated by the fraction of live cells of DMSO-treated control (counted using Trypan Blue staining and Countess II automated cell counter (Thermofisher). Data points represent mean±SD for one experiment, n=2, N=2. D, Quantification of the clonogenic survival of niraparib-treated OVCAR8 cells with CSNK2A2 CRISPR knock-out. Bars represent mean±SD for one experiment, n=2, N=1. E, Flow cytometry analysis of the EdU incorporation assay in MDA-MB-468 cells treated with silmitasertib and SGC-CK2-1 for 72 h. F, Quantification of the cell cycle phase distribution determined in EdU incorporation assay, related to Figure 3 A-D. G, Quantification of the micronuclei formation in COV318 and OVCAR8 cells treated with carboplatin for 24 h and then released to the carboplatin-free medium in presence or absence of 5 µM silmitasertib for 6 h. Confocal imaging of 500-1000 cells per condition. The bars represent mean±SD for one experiment, n=2, N=2. *P* values are calculated using a two-way ANOVA with Tukey post-test for multiple comparisons. *$0.01 < P \leq 0.05$, **$0.001 < P \leq 0.01$.

**Supplementary Figure S5.** A, Heat-map of the expression of RB pathway genes, CCLE gene expression dataset. B, WB for RB1 expression in OVCAR3-Cas9 cells infected with control guide RNA or sgRB1. C, Representative images of the clonogenic survival of sgRB1-transduced and control OVCAR3 cells treated with niraparib in presence or absence of silmitasertib. D, Quantification of the clonogenic assay in C.

**Supplementary Figure S6**. Duration of interphase in mitosis in individual cells quantified from the time-lapse videos SV1-16 taken at IncuCyte Zoom microscope using 10x objective for 96 h. Cells were exposed to niraparib for 48 h, followed by additional 48 h in presence or absence of silmitasertib. Rounding of the cells in prometaphase was considered the start of mitosis, while cell spreading after cytokinesis was considered as the end of the cell division. Detachment and signs of apoptosis (membrane blebbing) happening in mitosis were assigned as mitotic cell death. Exit from mitosis without division to 2 daughter cells was assigned as 4N. n = 50, N = 1.

**Supplementary Figure S7.** A, Quantification of the confocal imaging of the nuclear fluorescence intensity of the γH2AX immunostaining for the indicated cell lines treated for 72 h with silmitasertib. Cell line names in cerulean correspond to the RB1-deficient models where silmitasertib acted synergistically with carboplatin or niraparib; cell line names in violet correspond to the RB1-proficient models. Data points represent individual cells, and the red line represents the mean. B, Quantification of the nuclear fluorescence intensity of the γH2AX-Alexa647 or RPA-Alexa555 immunostaining for the indicated cell lines treated with 5 µM silmitasertib +/- 1 µM niraparib for 72 h. C, Flow cytometry-based analysis of the γH2AX immunostaining intensity in the different cell cycle phases after 72 h treatment with carboplatin +/- silmitasertib. The cell cycle phases were discriminated by EdU/propidium iodide staining as described for EdU incorporation assay in the Materials and methods. *P* values are calculated using a two-way ANOVA with Tukey post-test for multiple comparisons. *$0.01 < P \leq 0.05$, **$0.001 < P \leq 0.01$, ***$0.0001 < P \leq 0.001$ and ****$P \leq 0.0001$.

**Supplementary Figure S8.** A, Immunoblotting analysis of the replicative DNA damage signaling triggered by replication blocking nucleotide depletion by hydroxyurea in presence or absence of CK2 inhibitor. Cells were pre-treated for 24 with silmitasertib to achieve full loss of CK2 activity, and then were treated with acute dose of HU for 45 min as described[36]. B, Quantification of the imaging for immunostained inter-strand crosslink repair foci of FANCD2 in the pulse-labeled EdU-positive cells after 72 h of silmitasertib treatment. C, Quantification of the nuclear fluorescence intensity of the γH2AX-Alexa647 or RPA-Alexa555 immunostaining for the indicated cell lines treated with silmitasertib and HU as described in A, or with the combination of silmitasertib and niraparib for 72 h. Dots represent individual nuclei, 500-2000 nuclei analyzed by confocal imaging. Cell line names in cerulean correspond to the RB1-deficient models where silmitasertib acted synergistically with carboplatin or niraparib; cell line names in violet correspond to the RB1-proficient models. *P* values are calculated using a two-way ANOVA with Tukey post-test for multiple comparisons. *$0.01 < P \leq 0.05$, **$0.001 < P \leq 0.01$, ***$0.0001 < P \leq 0.001$ and ****$P \leq 0.0001$.

**Supplementary Figure S9.** A, Toxicity of niraparib in OVCAR3 and OVCAR4 cells transfected with 2 pooled siRNAs against BRCA2 or non-targeting control. The fraction of dead cells was calculated from the imaging analysis of the cells stained with Hoechst and CellTox Green toxicity dye after 72 h of treatment with the indicated drugs. B, Quantification of the micronuclei formation in the cells treated as in A. 200-500 cells were quantified per replica, n=3. C, Quantification of the imaging for immunostained DNA repair foci of RAD51 and pS1524 BRCA1 performed to evaluate the efficacy of the RAD51 recombinase inhibitor B02, further used in D. Confocal imaging of 200-500 nuclei, *P* values are calculated using a two-way ANOVA with Tukey post-test for multiple comparisons. *$0.01 < P \leq 0.05$, **$0.001 < P \leq 0.01$, ***$0.0001 < P \leq 0.001$ and ****$P \leq 0.0001$. D, Clonogenic survival of OVCAR3 and COV318 exposed to carboplatin or combination of carboplatin and silmitasertib in presence of RAD51 inhibitor B02 for 72 h, followed by recovery in drug-free medium for 10 days. Cell line names in violet mark the RB1-proficient models.

**Supplementary Figure S10**

A-C, Confocal imaging of RB1-deficient EOC172, EOC227, and EOC883p organoids treated with niraparib and silmitasertib for 7 days. Hoechst and CellTox Green staining for 6 h at 37°C.

### Supplementary figure S1

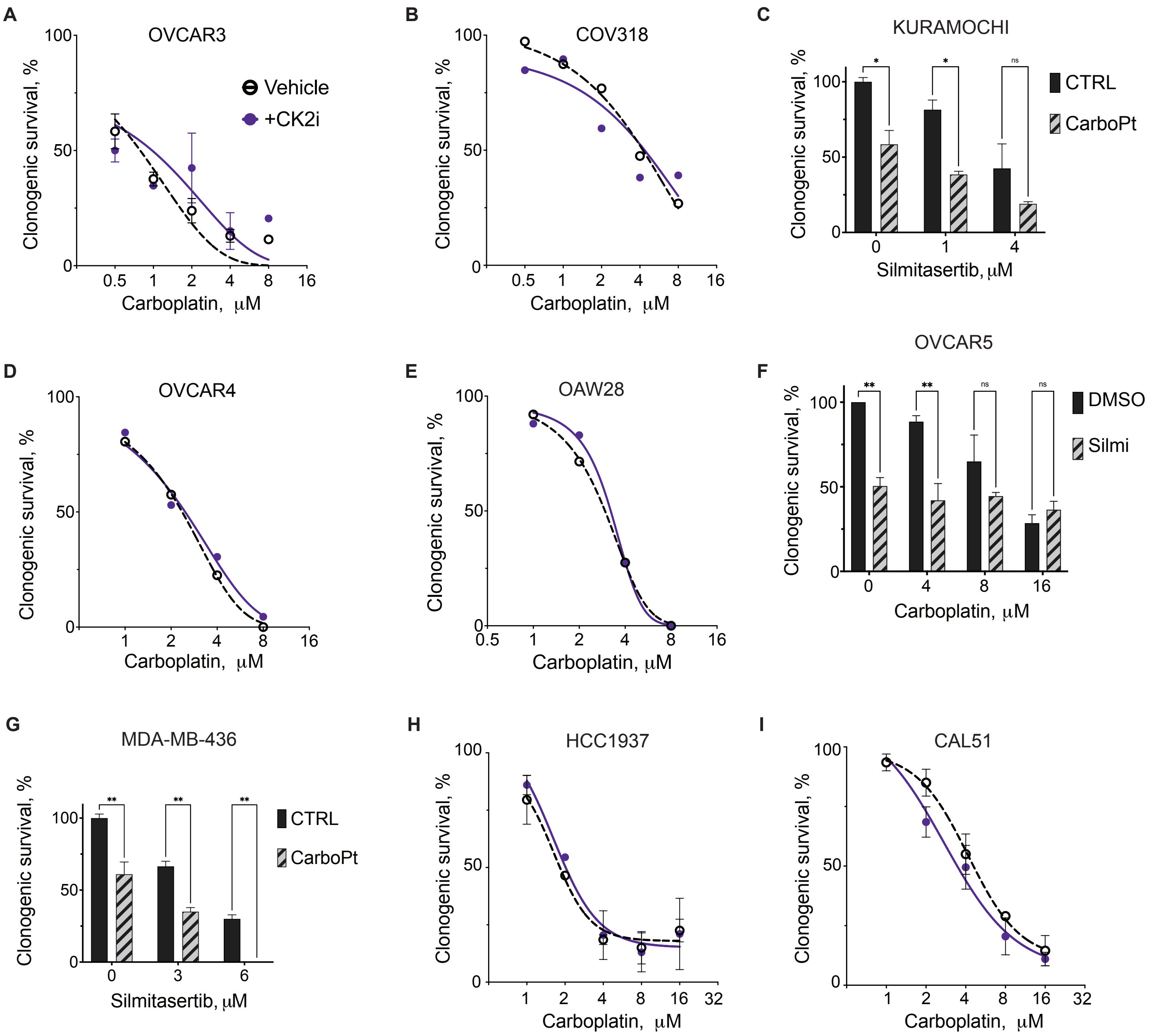

#### Supplementary Figure S2

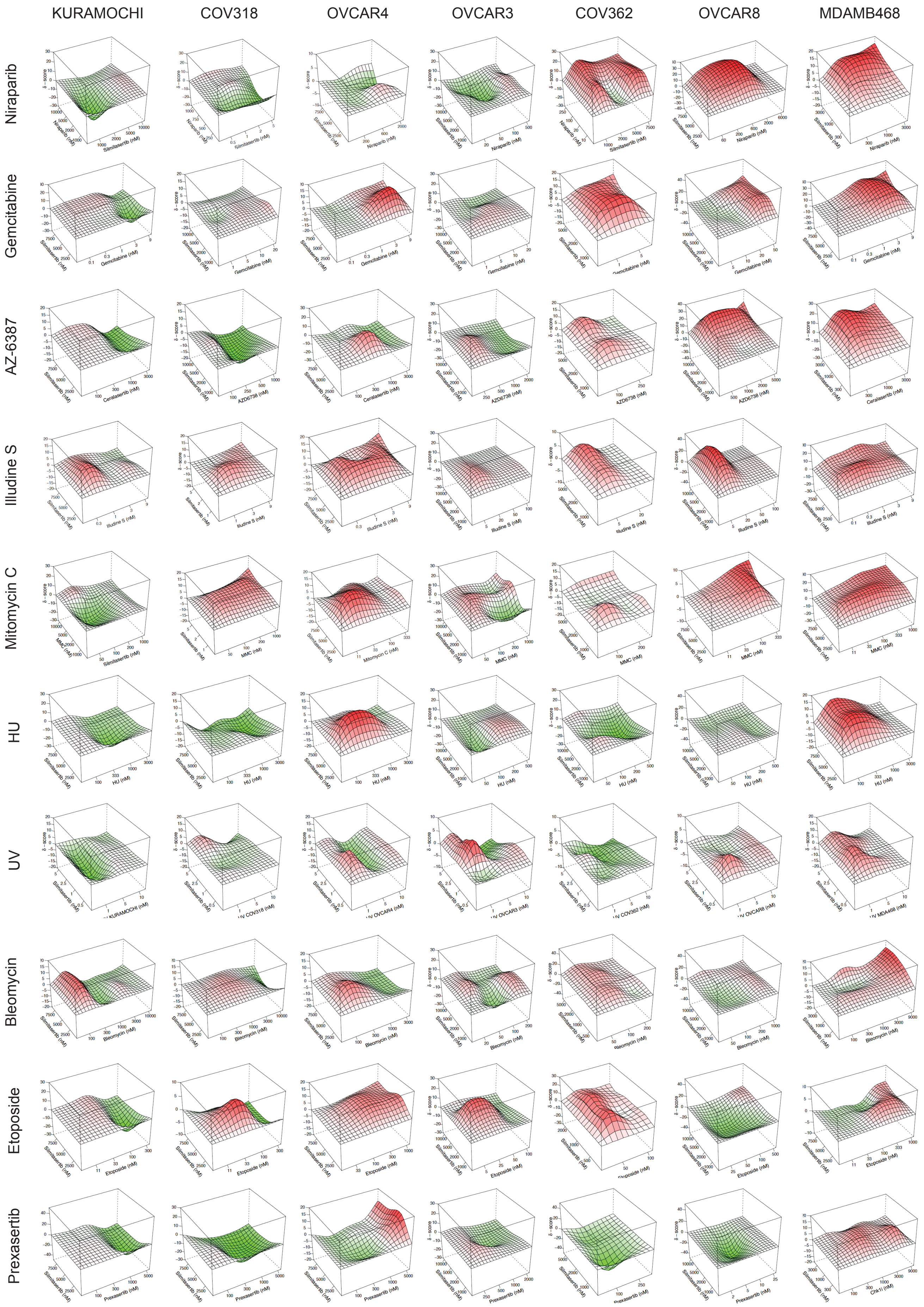

**Supplementary Figure S3**

A

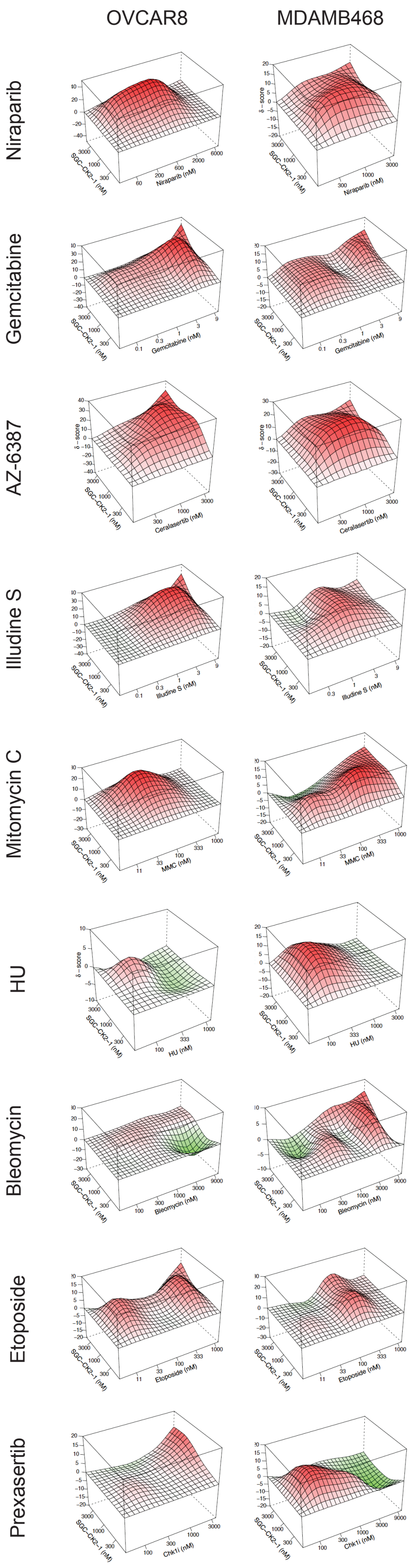


B

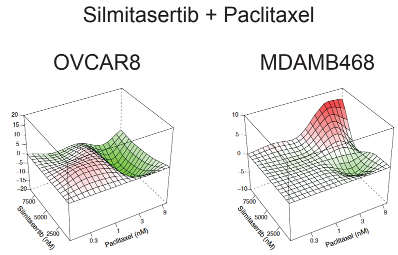

**Supplemenbtary Figure S4**

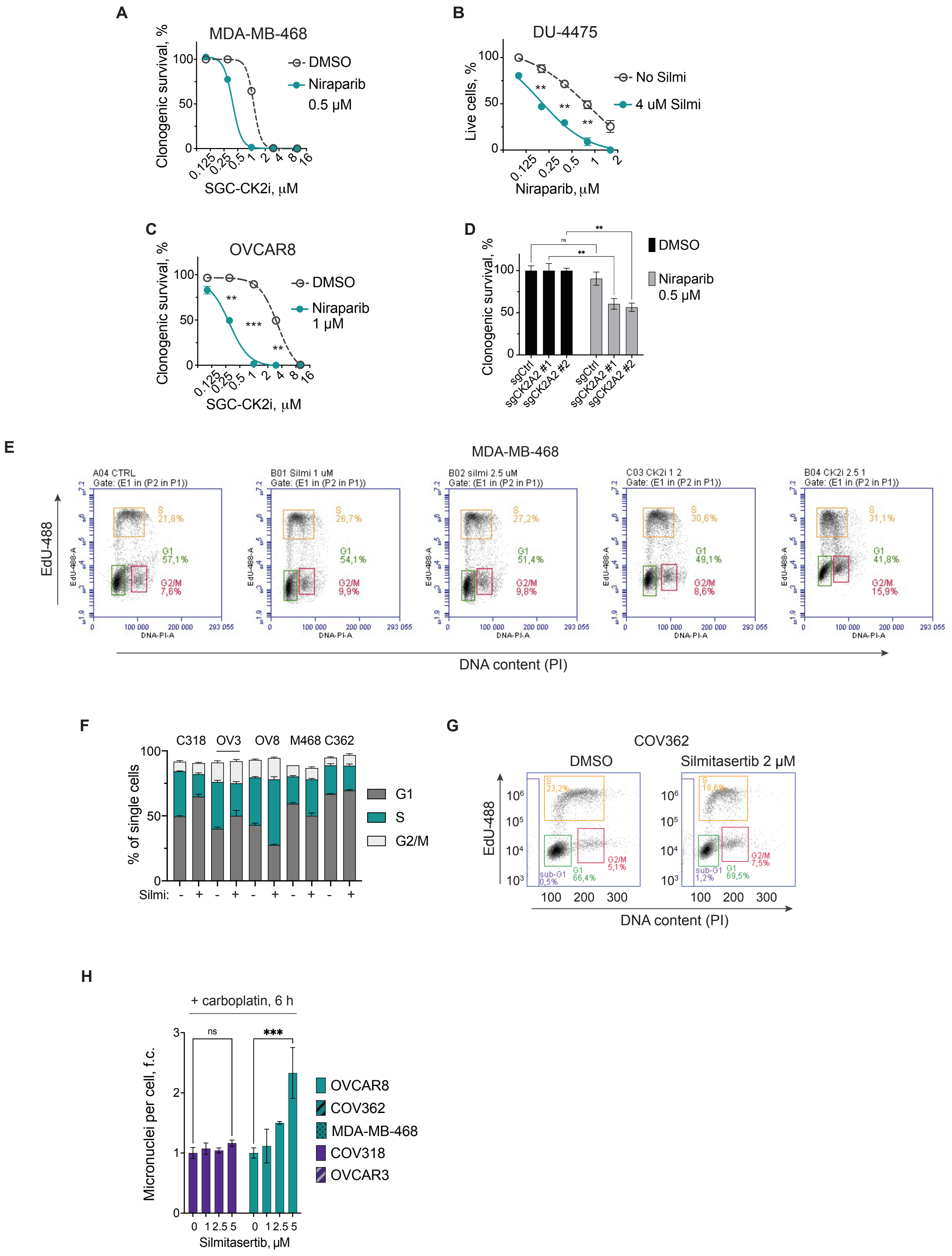

#### Supplementary Figure S5

A

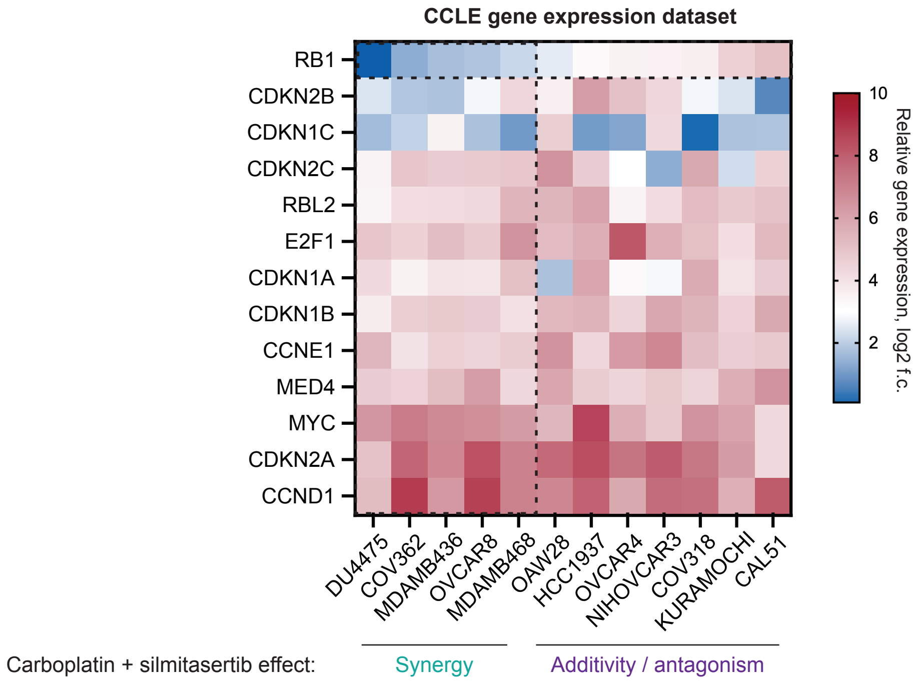


B

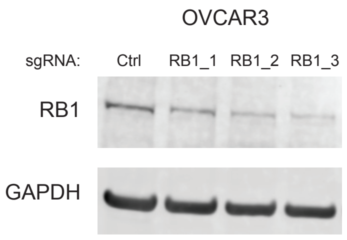


C

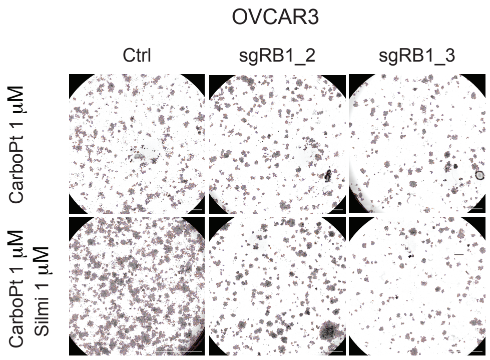


D

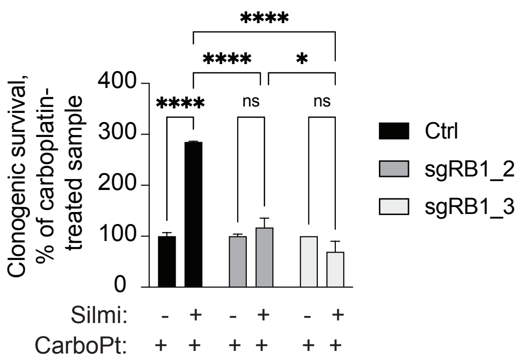

**Supplementary Figure S6**

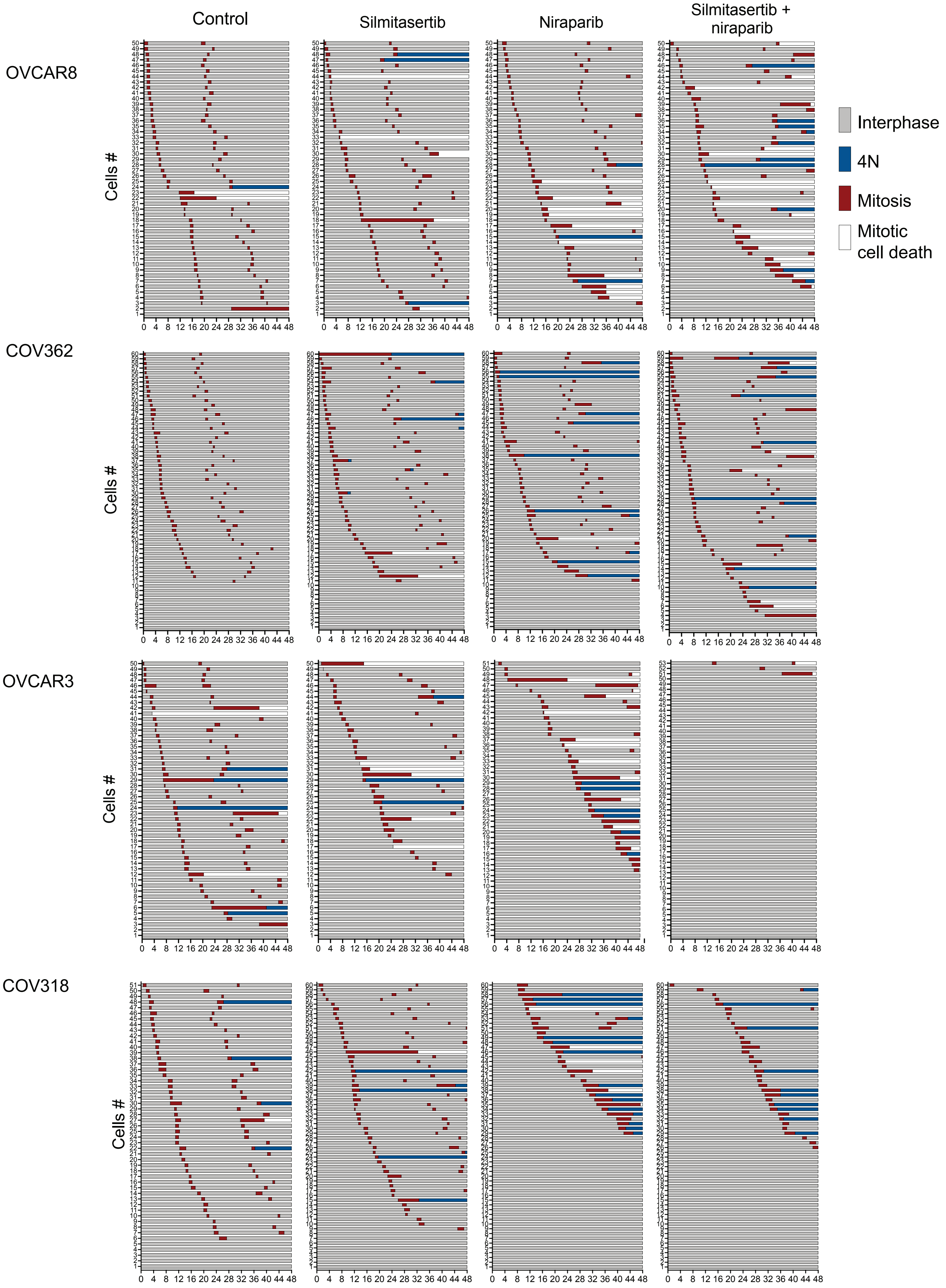

**Supplementary Figure S7**

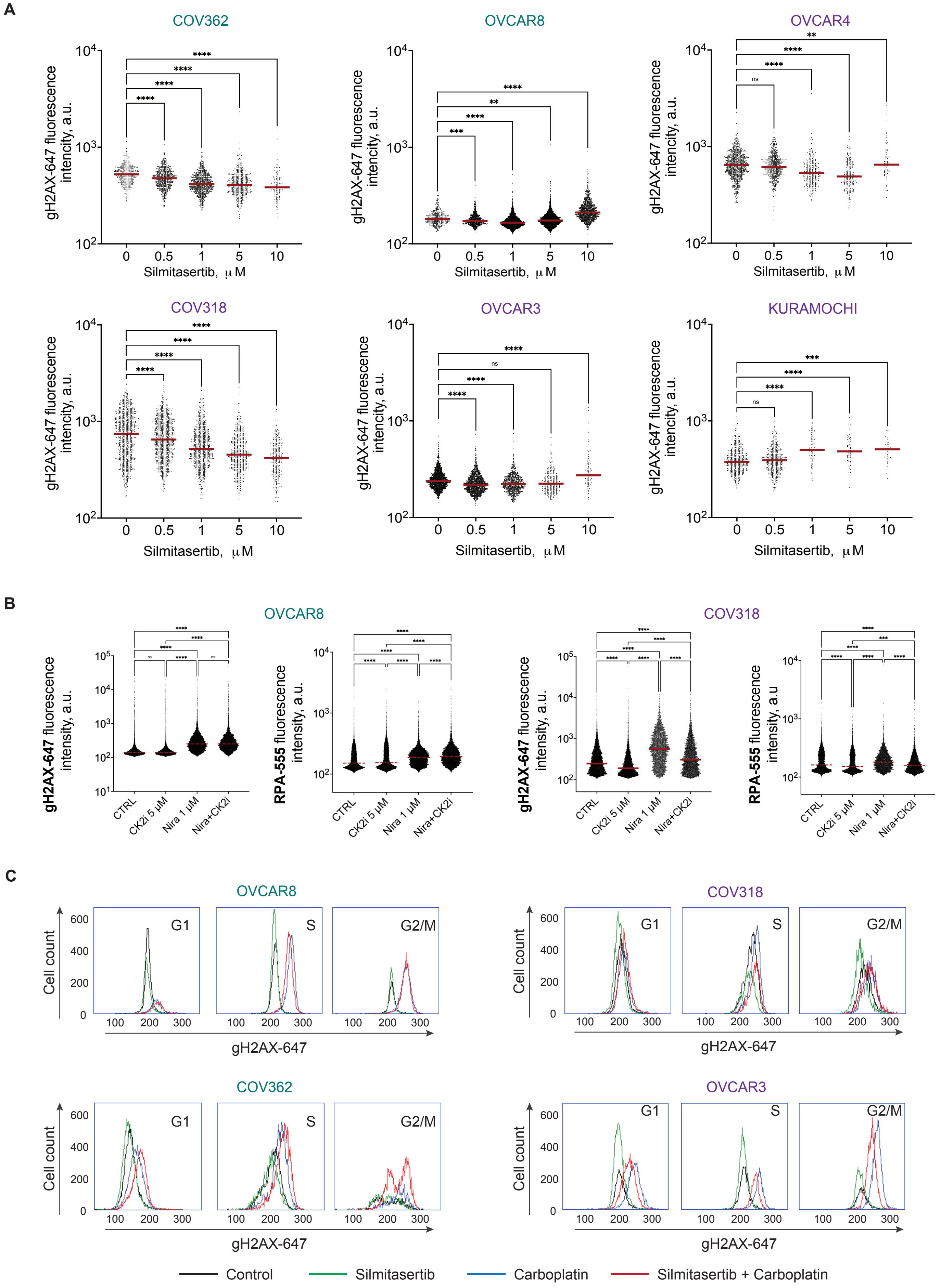

**Supplementary Figure S8**

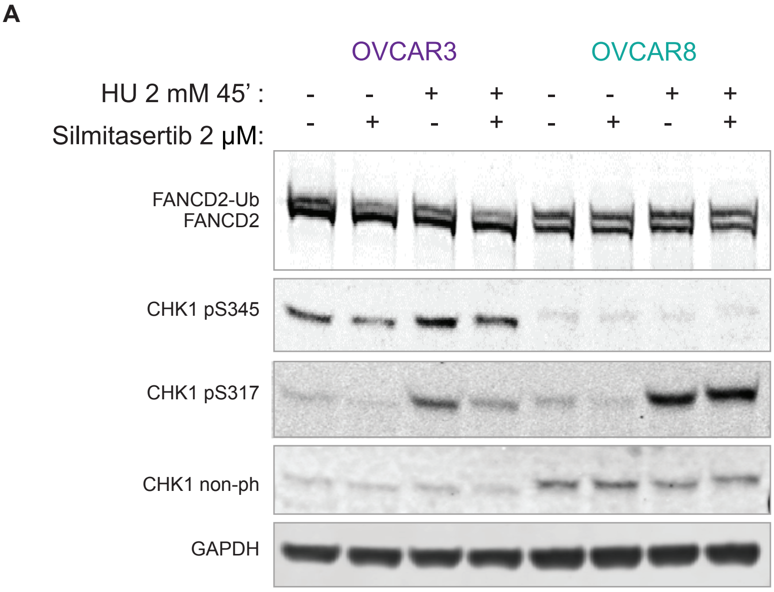


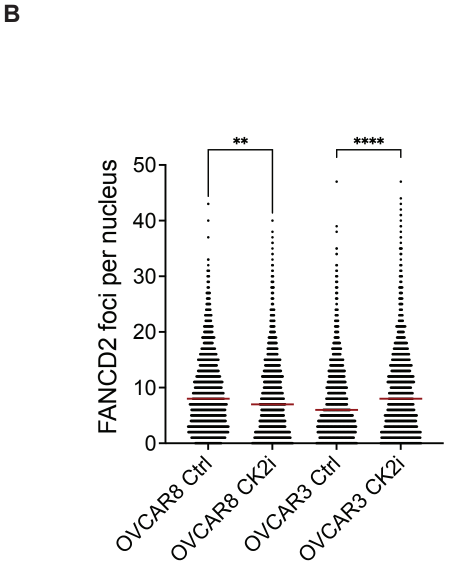


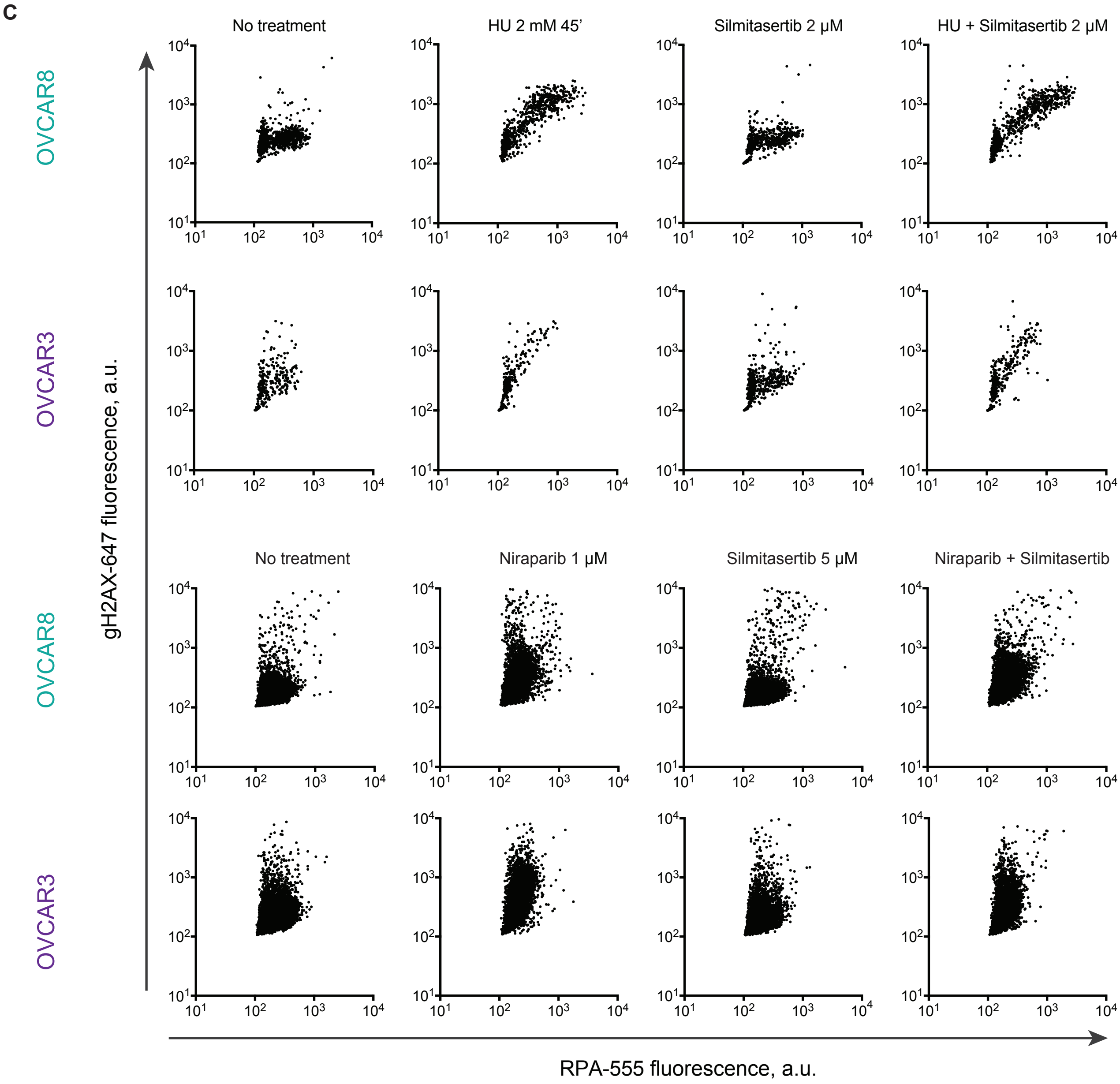

**Supplementary Figure S9**

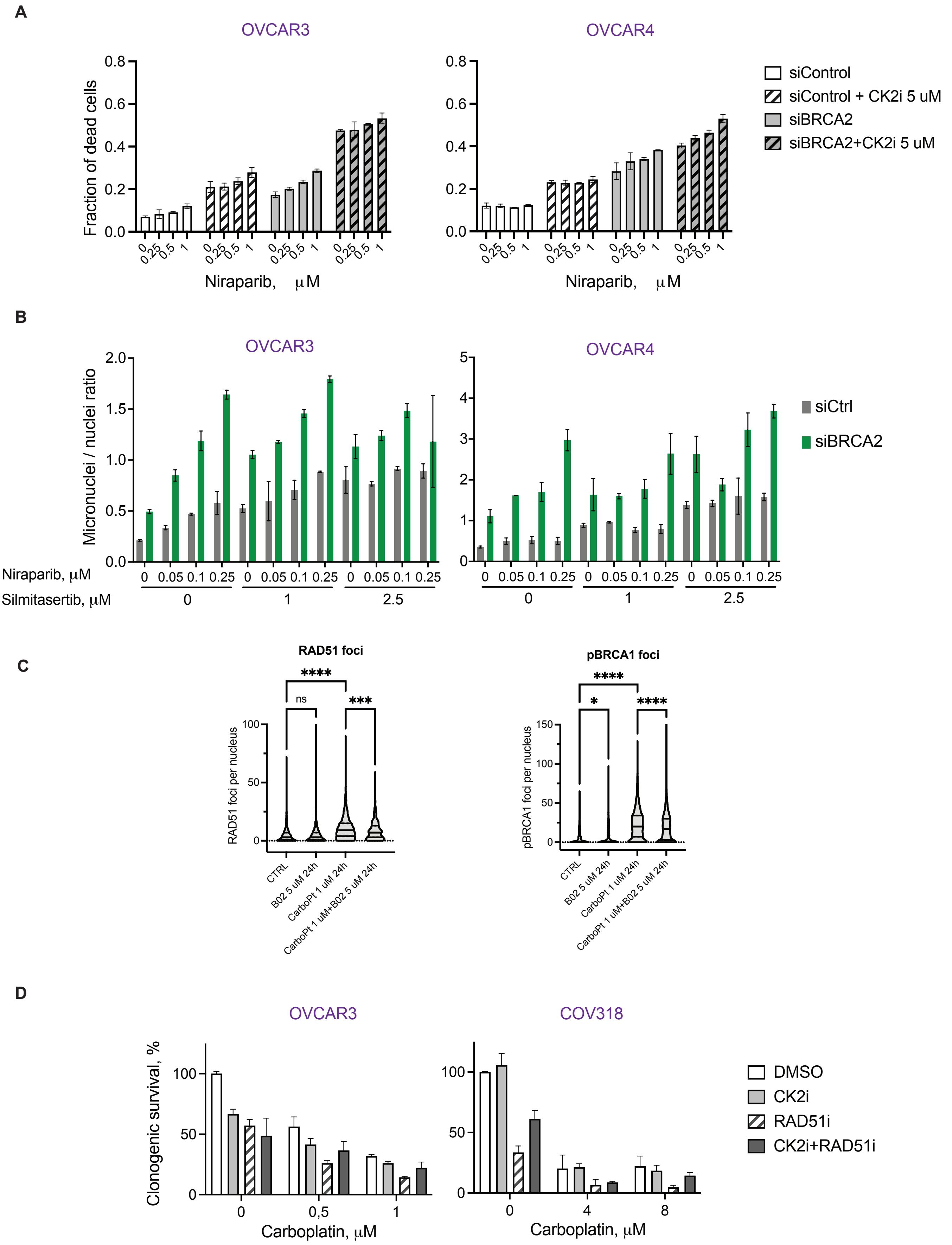

**Supplementary Figure S10**

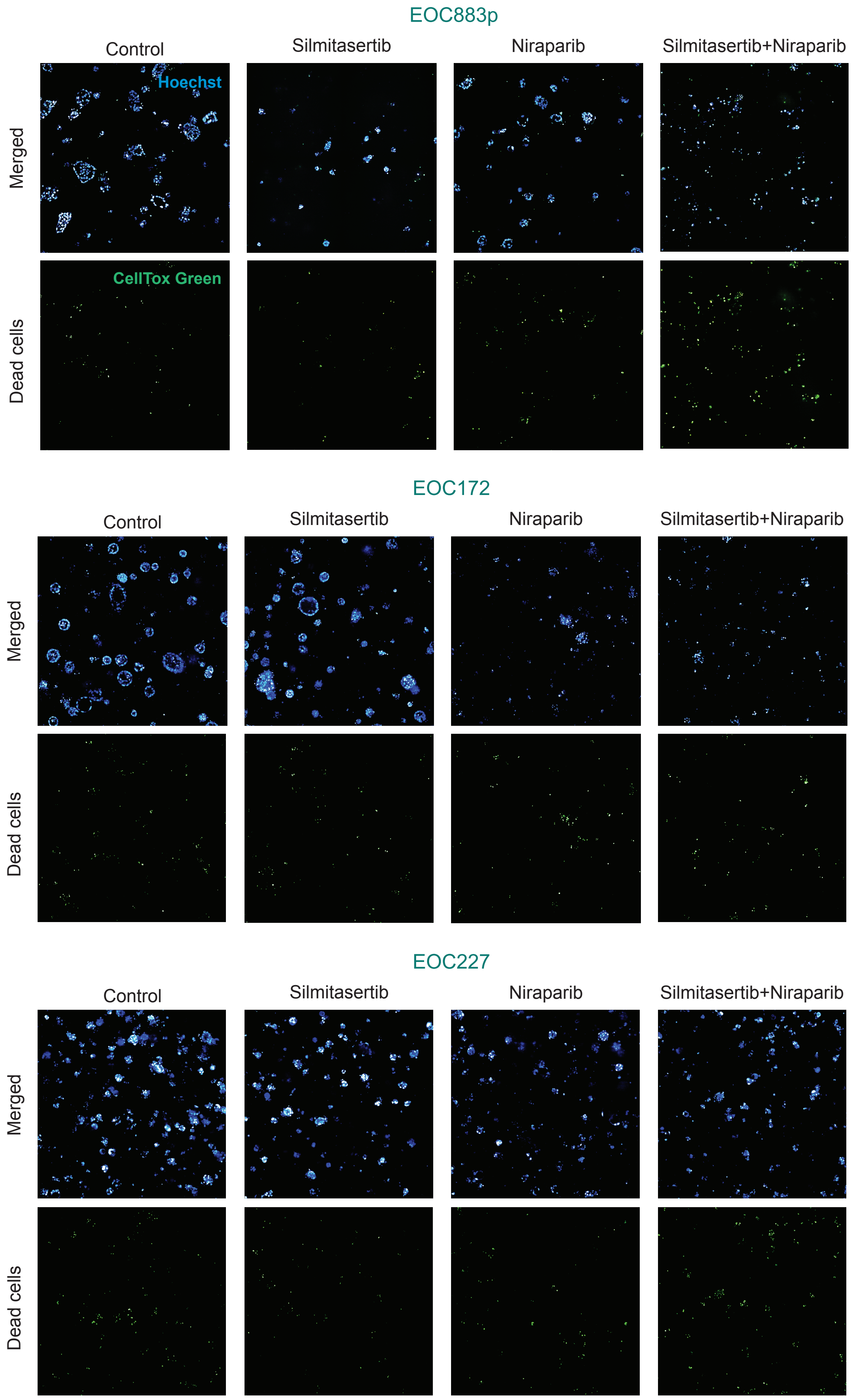
